# LY-CoV555, a rapidly isolated potent neutralizing antibody, provides protection in a non-human primate model of SARS-CoV-2 infection

**DOI:** 10.1101/2020.09.30.318972

**Authors:** Bryan E. Jones, Patricia L. Brown-Augsburger, Kizzmekia S. Corbett, Kathryn Westendorf, Julian Davies, Thomas P. Cujec, Christopher M. Wiethoff, Jamie L. Blackbourne, Beverly A. Heinz, Denisa Foster, Richard E. Higgs, Deepa Balasubramaniam, Lingshu Wang, Roza Bidshahri, Lucas Kraft, Yuri Hwang, Stefanie Žentelis, Kevin R. Jepson, Rodrigo Goya, Maia A. Smith, David W. Collins, Samuel J. Hinshaw, Sean A. Tycho, Davide Pellacani, Ping Xiang, Krithika Muthuraman, Solmaz Sobhanifar, Marissa H. Piper, Franz J. Triana, Jorg Hendle, Anna Pustilnik, Andrew C. Adams, Shawn J. Berens, Ralph S. Baric, David R. Martinez, Robert W. Cross, Thomas W. Geisbert, Viktoriya Borisevich, Olubukola Abiona, Hayley M. Belli, Maren de Vries, Adil Mohamed, Meike Dittmann, Marie Samanovic, Mark J. Mulligan, Jory A. Goldsmith, Ching-Lin Hsieh, Nicole V. Johnson, Daniel Wrapp, Jason S. McLellan, Bryan C. Barnhart, Barney S. Graham, John R. Mascola, Carl L. Hansen, Ester Falconer

## Abstract

SARS-CoV-2 poses a public health threat for which therapeutic agents are urgently needed. Herein, we report that high-throughput microfluidic screening of antigen-specific B-cells led to the identification of LY-CoV555, a potent anti-spike neutralizing antibody from a convalescent COVID-19 patient. Biochemical, structural, and functional characterization revealed high-affinity binding to the receptor-binding domain, ACE2 binding inhibition, and potent neutralizing activity. In a rhesus macaque challenge model, prophylaxis doses as low as 2.5 mg/kg reduced viral replication in the upper and lower respiratory tract. These data demonstrate that high-throughput screening can lead to the identification of a potent antiviral antibody that protects against SARS-CoV-2 infection.

**One Sentence Summary:** LY-CoV555, an anti-spike antibody derived from a convalescent COVID-19 patient, potently neutralizes SARS-CoV-2 and protects the upper and lower airways of non-human primates against SARS-CoV-2 infection.

---

The global COVID-19 pandemic continues to spread rapidly with substantial health, economic, and societal impact.(*1*) Severe acute respiratory syndrome coronavirus-2 (SARS-CoV-2), the novel coronavirus responsible for COVID-19 disease, can induce acute respiratory distress syndrome and a wide spectrum of symptoms leading to substantial morbidity and mortality.(*2*) Neutralizing antibodies represent an important class of therapeutics which could provide immediate benefit in treatment or as passive prophylaxis until vaccines are widely available. Passive prophylaxis could be an alternative to vaccination in populations where vaccines have been found to be less efficacious.(*3, 4*)

SARS-CoV-2 neutralizing antibody discovery efforts have focused on targeting the multi-domain surface spike protein, a trimeric class I fusion protein that mediates viral entry. Spike protein-dependent viral entry is initiated by upward movement of the receptor-binding domain (RBD) at the apex of the protein allowing access to bind the angiotensin converting enzyme 2 (ACE2) cellular receptor.(*5–8*) Upon receptor engagement, coordinated proteolytic cleavage, shedding of the S1 subunit, and conformational rearrangement of the S2 subunit, leads to viral fusion with the cell and transfer of genetic material. Given the critical nature of the RBD interaction with ACE2 for viral entry, antibodies that bind the RBD and interfere with ACE2 binding can have potent neutralizing activity.(*9–11*)

To test the potential for neutralizing monoclonal antibodies (mAbs) to prevent SARS-CoV-2 infection *in vivo*, we used the rhesus macaque challenge model. While rhesus macaques do not exhibit the severe pulmonary symptoms sometimes associated with human COVID-19 disease, the model allows assessment of viral replication in the upper and lower airways.(*12–16*) Of particular interest, recent studies in this model have shown that prior exposure to SARS-CoV-2 or administration of a SARS-CoV-2 vaccine are sufficient to prevent infection upon subsequent challenge.(*15, 17*) Protecting non-human primates (NHPs) from SARS-CoV-2 infection may inform the clinical development of medical countermeasures for COVID-19.(*14, 18*)

In this study, we report a strategy for high-throughput screening, which allowed for the rapid identification and subsequent characterization of anti-spike neutralizing antibodies. An RBD-specific antibody (LY-CoV555) was discovered that can bind the RBD in the up (active) or down (resting) conformation and potently neutralize SARS-CoV-2 *in vitro*. Passive immunization by infusion of LY-CoV555 protected both upper and lower airways from SARS-CoV-2 infection in a rhesus macaque model. These data supported the progression of LY-CoV555 into clinical evaluation.

## Identification and characterization of SARS-CoV-2 neutralizing antibodies

To identify potential therapeutic antibodies from a convalescent COVID-19 patient, a novel high-throughput screening approach was used to identify relevant anti-spike mAbs (Fig. 1, Fig. S1A). Peripheral blood mononuclear cells (PBMCs) were obtained approximately 20 days post onset of symptoms. Two screening assays were utilized: (1) a multiplexed bead-based assay using optically-encoded microbeads, each conjugated to either soluble prefusion-stabilized trimeric SARS-CoV-2 or SARS-CoV spike protein and (2) a live cell-based assay using mammalian cells that transiently express full-length membrane-anchored SARS-CoV-2 spike protein (Fig. S1A). In total, 5.8 million PBMCs were screened over three days, and machine learning (ML)-based analysis pipelines were used to automatically select and rank > 4,500 antibody “hits” (0.08% frequency), of which 2,238 single antibody-secreting cells were chosen for recovery. Next-generation sequencing libraries of antibody genes from selected single B-cells were generated and sequenced, and a custom bioinformatics pipeline with ML-based sequence curation was used to identify paired-chain antibody sequences, resulting in 440 unique high-confidence paired heavy and light chain sequences (Fig. 1). The sequences belonged to 394 clonal families and used a diverse set of 39 VH genes, with the VH3 family of genes representing 57% of total diversity (Fig. S1B), similar to other reports.(*19*) Among these, the VH3-30 gene was the most common (39%) (Fig. S1B). Of the 440 unique antibodies identified, 4% were cross-reactive to both full-length SARS-CoV-2 and SARS-CoV spike proteins. The mean sequence identity to germline was high (98% and 99% for heavy and light chains respectively; Fig. S1B), likely due to sample collection early in the immune response.

**Fig. 1.**
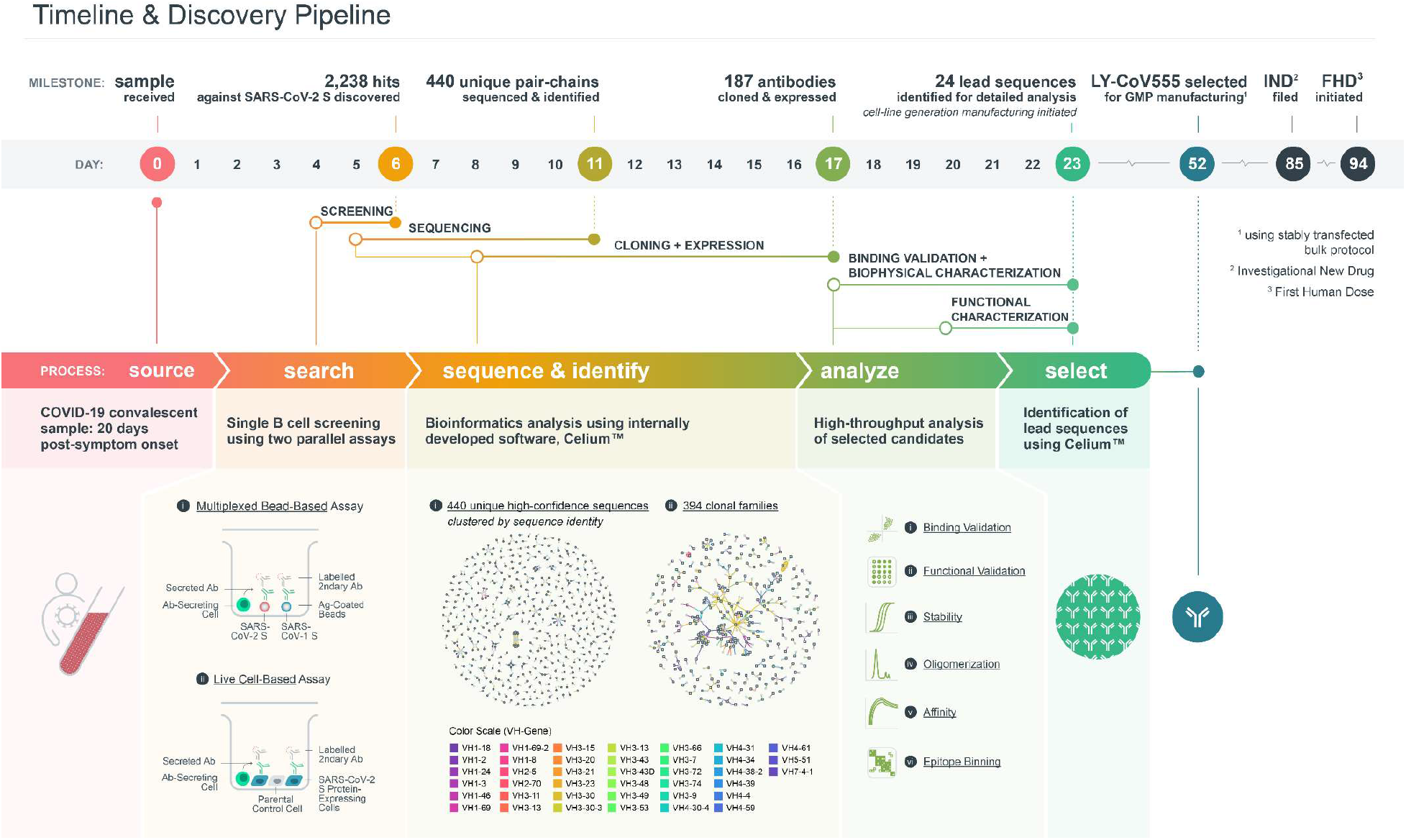
Timeline, screening and sequence analysis. (Top) Timeline of events leading to clinical evaluation of LY-CoV555. (Bottom) Outline of the discovery process. (Left) PBMC collection from a COVID-19 patient and antibody search using multiplexed bead-based and live cell-based screening assays based on structurally-defined and stabilized spike protein probes. In the multiplexed assay, beads were conjugated to either SARS-CoV-2 or SARS-CoV spike protein. In the live-cell assay, parental cells were visualized by a passive dye, and SARS-CoV-2 spike protein-expressing cells by GFP fluorescence. Positive binding was detected by fluorescently labeled anti-human IgG secondary antibodies. (Middle) Sequence analysis of the 440 unique high-confidence paired-chain antibodies. Graphical representation of antibodies clustered according to sequence identity (Middle Left) or clonal family relationships (Middle Right). Each node indicates a chain (heavy or light), or a cluster of chains (heavy or light). E ach line indicates a single antibody, colored by VH gene usage. Multiple lines that connect to the same heavy and light chain clusters represent clonally related antibodies. (Right) Representation of the high-throughput analyses utilizing binding and functional validation for lead antibody candidate selections. GFP = green fluorescent protein; IgG = immunoglobulin G.

From the set of 440 antibodies, we used an internally developed informatics and data visualization software package, Celium™, to select 187 antibodies for rapid cloning and expression. Preference was given to antibodies observed at high frequency across the dataset, especially those discovered in both multiplexed soluble protein and live-cell assays. The selection also maximized the diversity of VH genes and CDR3 sequences, and limited CDR3 sequence liabilities. A total of 175 sequences were successfully cloned into expression vectors to generate recombinant antibodies with immunoglobulin G1 (IgG1) backbones for more detailed characterization. Subsequent characterization included high-throughput biophysical analysis (Fig. S2A), validation of soluble and cell-associated spike protein binding, cross reactivity to other coronavirus spike proteins and three circulating SARS-CoV-2 spike variants (Fig. S2B), apparent binding affinity to soluble spike by surface plasmon resonance (SPR) (Fig. 2A), and functional screening in a high-throughput pseudotyped lentivirus reporter neutralization assay (Fig. 2C).

**Fig. 2.**
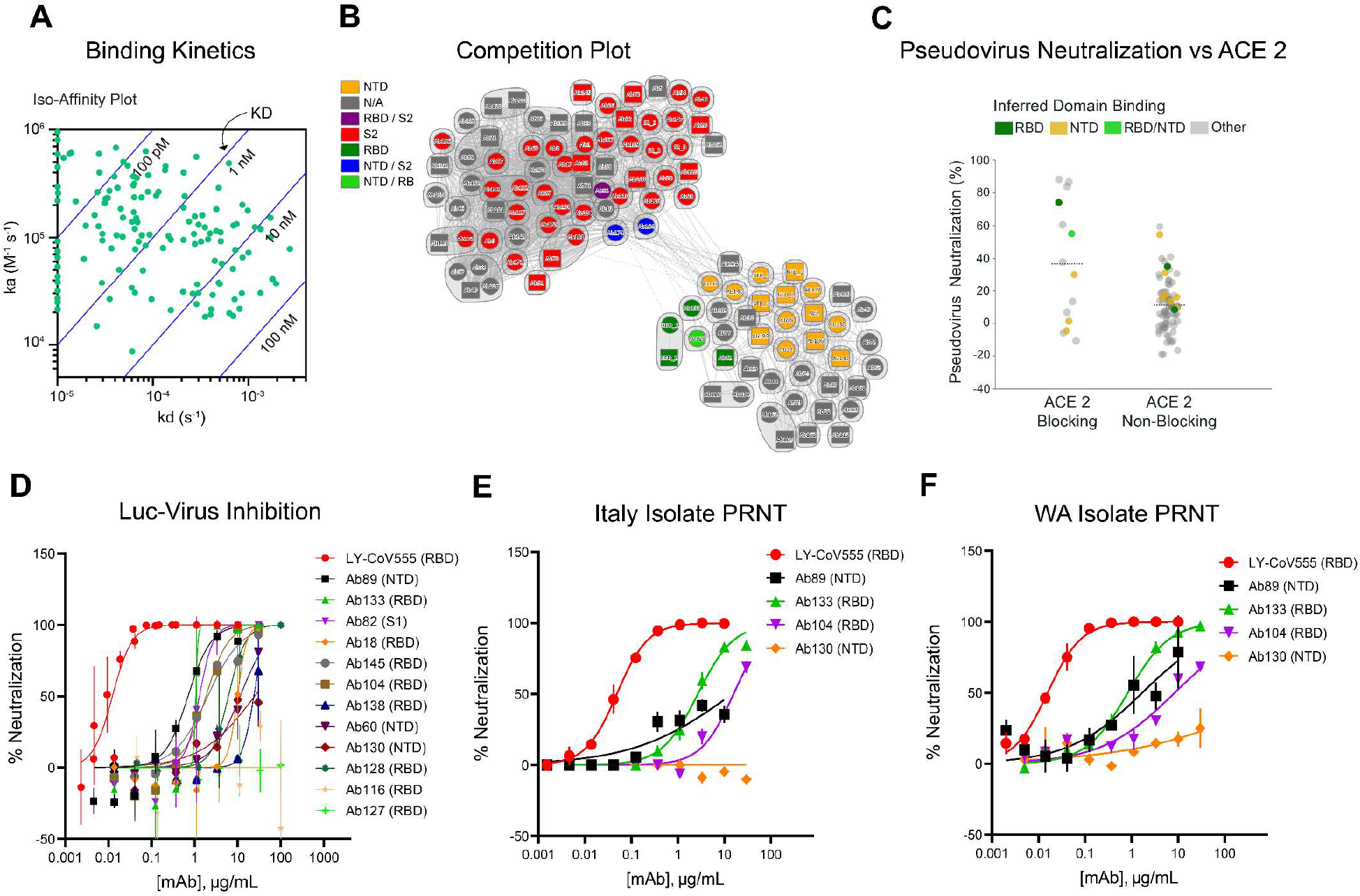
Binding, epitope pairing, and functional characterization of discovered antibodies. (A) Binding kinetics of recombinantly expressed antibodies. Association and dissociation rate constants were measured by high-throughput surface plasmon resonance (SPR) capture kinetic experiments with antibodies as immobilized ligands and antigens of interest as analytes. The distribution of kinetic values is displayed in an iso-affinity plot. (B) Competition plot of recombinantly expressed antibodies. Each antibody was tested in two orientations: as a ligand on the chip, and as an analyte in solution. Individual antibodies are represented either as a circle (data present in both orientations) or as a square (data present with the antibody in a single orientation). Bins are represented as envelopes (95 total) and competition between antibodies as solid (symmetric competition) or dashed (asymmetric competition) lines. Benchmark-based blocking profiles are indicated by color. Based on benchmark competition profiles, a clear divide between S1- and S2-specific antibodies is seen with low meshing between the groups. (C) Recombinantly expressed antibodies were screened utilizing a high-throughput pseudotyped lentivirus reporter neutralization assay (antibody blocking profiles are indicated by color and delineated by ACE2 competition). (D) Neutralization of recombinant SARS-CoV-2 encoding nanoluciferase in the Orf7a/b locus in infected Vero-E6 cells 24 hours post inoculation (values plotted are means of two replicates, with error bars showing SEM). Plaque reduction neutralization test (PRNT) assay for Italian INMI-1 isolate (E) and 2020/USA/WA1 isolate (F) of SARS-CoV-2 in Vero-E6 cells 72 hours post inoculation; values plotted are means of two replicates, with error bars showing SEM. SEM = standard error of mean.

High-throughput SPR experiments were used to characterize the epitope coverage of the 175 antibodies: these experiments included antibody pairing, isolated domain binding, and binding competition with ACE2 (Fig. 2). Benchmark antibodies with known binding to S1 subunit, N-terminal domain, RBD, and S2 subunit epitopes of the SARS-CoV spike protein and cross-reactivity to SARS-CoV-2 spike protein were included to mark epitope identity. Antibody cross-blocking results are summarized in the competition plot (Fig. 2B), as well as in the heat map (Fig. S3). In total, 95 unique bins (including controls) were identified, and a clear divide between S1- and S2-specific antibodies as inferred by benchmark competition was seen (Fig. 2B, Fig. S3), suggesting that these antibodies possessed a broad epitope diversity. Only approximately 10% of the antibodies tested exhibited ACE2 competition. Antibodies with ACE2 binding inhibition properties had the greatest neutralizing activity based on pseudotyped lentivirus reporter neutralization (Fig. 2C), although antibodies to other domains also had detectable neutralizing activity.

A lead panel of 24 antibodies (Table S1) was selected using the Celium™ software, based on the following criteria: (1) binding to SARS-CoV-2 spike protein in either the multiplexed bead-based or the live cell-based validation assay, (2) >30% pseudovirus neutralizing activity at any of the concentrations tested (10, 1, 0.1, or 0.01 μg/mL), (3) dose-dependent neutralization profile, (4) RBD competition, (5) ACE2 blocking activity, and (6) acceptable biophysical profile (melting temperature, solubility, and polydispersity). The selected antibodies were then produced at larger scale for further functional testing, epitope mapping, and structural analysis (summarized in Table S2, Table S3, Fig. S4, Fig. S5).

The selected antibodies had a broad range of neutralizing activity in multiple *in vitro* assays including pseudovirus (Table S4) and various full virus assay formats (Fig. 2). Using a replication-competent SARS-CoV-2 molecular clone modified with a nano-luciferase reporter virus, (Fig. 2D), half maximal inhibitory concentration (IC_50_) neutralizing activity values spanning nearly three orders of magnitude were observed (Table S4). For a smaller number of antibodies, viral neutralization was further characterized in a Plaque Reduction Neutralization Test (PRNT) format against two different clinical SARS-CoV-2 isolates (Fig. 2 E, F), the Italian INMI-1 isolate (clade 19A) and the USA/Wa-1/2020 isolate (clade 19B), representing the two major clades of SARS-CoV-2 (www.gisaid.com). Interestingly, it was observed that some non-RBD binding antibodies, for example Ab82, Ab89, and Ab130, exhibited greater neutralizing activity in some of the live virus SARS-CoV-2 assays compared to pseudovirus assays (Table S4). Notably, the neutralization potency of one mAb, Ab169 (designated LY-CoV555), an RBD binder and ACE2 blocker, was consistently and substantially greater than the rest and was selected for further development.

LY-CoV555 possessed greater neutralization potency relative to other identified RBD-binding and ACE2-blocking antibodies (e.g. Ab128 and Ab133), despite similar apparent binding affinities (Table S2), suggesting a distinct binding mode of recognition. Structural analysis using X-ray crystallography and cryo-electron microscopy (cryo-EM) demonstrated that two of the RBD-binding mAbs (Ab128 and Ab133) bind in a nearly identical fashion to one another (Fig. S5), and in a similar fashion to the previously described mAb CB6.(*10*) LY-CoV555 was observed to bind to an epitope overlapping the ACE2 binding site (Fig. 3A, B, and C); specifically, 7 of the approximate 25 sidechains in the RBD observed to form contact with ACE2.(*5, 20, 21*) Based on the crystal structure, the LY-CoV555 epitope was predicted to be fully accessible on both the up and down conformations of the RBD. This was confirmed by high resolution cryo-EM imaging of LY-CoV555 Fab complexes in which the LY-CoV555 Fab was observed to bind the spike protein RBD in both up and down conformations (Fig. 3D). This unique property is analogous to the binding of the Ebola mAb114 that binds the GP receptor-binding domain in both the pre-activation and activated states.(*22*) mAb114 was subsequently shown to effectively treat Ebola disease as monotherapy.(*23*) This may indicate an advantage for mAbs that can bind critical functional domains of class I fusion proteins at multiple stages of the entry process.

**Fig. 3.**
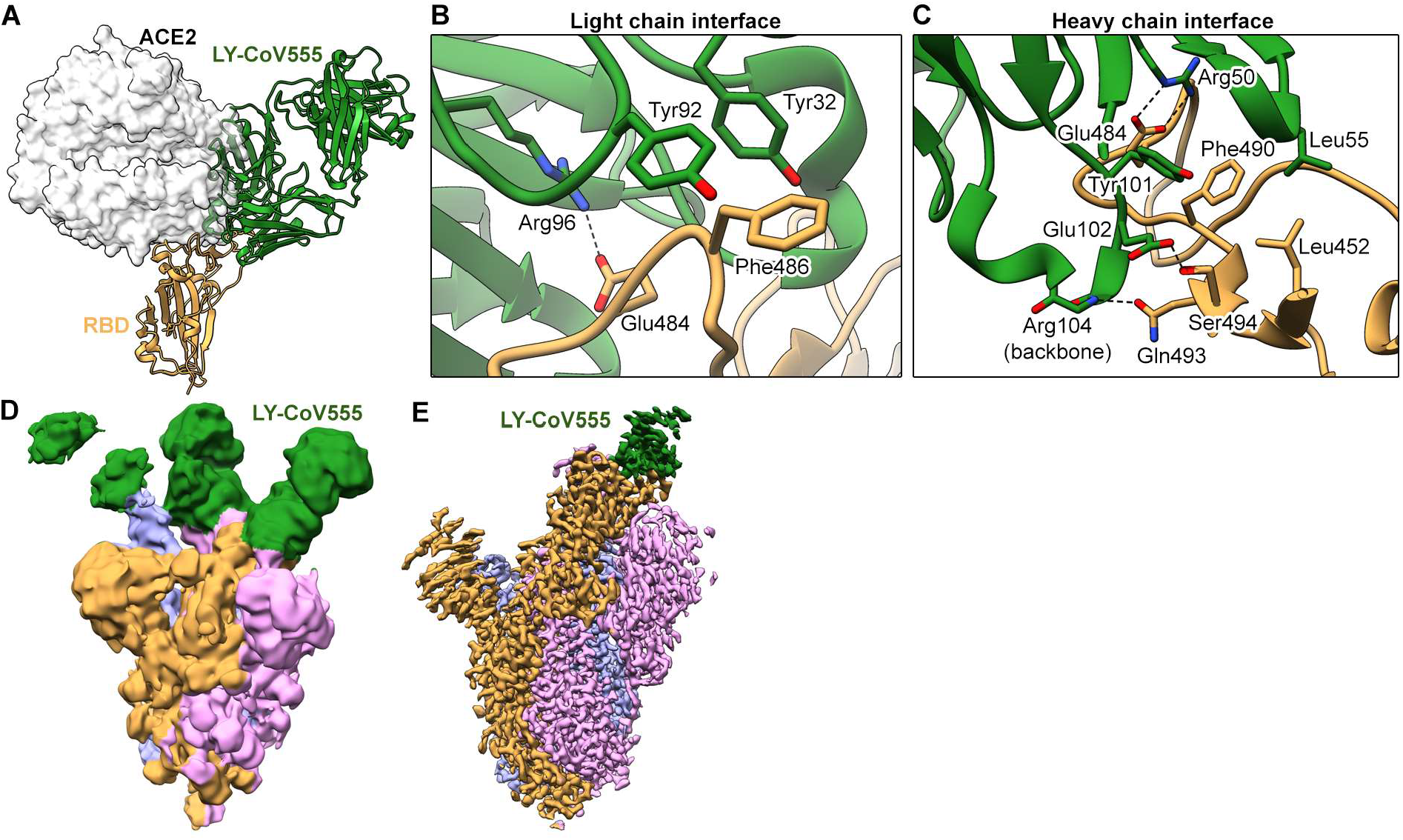
LY-CoV555 blocks ACE2 and binds to the spike protein RBD in the up and down conformations. (A) Crystal structure of the RBD-LY-CoV555 complex superimposed with the ACE2 receptor from a structure of the RBD-ACE2 complex (PDB ID: 6M0J) (*26*). Zoomed-in view of key atomic interactions at the interface of the LY-CoV555 light chain (B) and heavy chain (C) with the spike RBD. (D) Cryo-EM structure of the LY-CoV555-spike complex low-pass filtered to 8Å resolution and shown at low threshold in order to visualize all 3 Fabs (shown in green). (E) High-resolution cryo-EM map of the LY-CoV555-spike complex. Cryo-EM = cryo-electron microscopy; RBD = receptor-binding domain.

## LY-CoV555 provides protection from infection and viral replication in an NHP model of SARS-CoV-2 infection

To assess the ability of LY-CoV555 to protect from viral challenge, rhesus macaques were dosed intravenously (IV) with 1, 2.5, 15 or 50 mg/kg of LY-CoV555 or 50 mg/kg of a control IgG1 antibody 24 hours prior to virus challenge. LY-CoV555 doses were chosen to provide a range of serum antibody concentrations and inform subsequent clinical dosing. Macaques were inoculated intranasally and intratracheally with a total of 1.1×10^5^ pfu of SARS-CoV-2 (USA-WA1/2020) and were monitored by twice daily cage-side observations and respiratory exams throughout the study. Respiratory and clinical signs of disease in the macaques were limited, and generally mild lobar congestion and hyperemia were observed macroscopically across control and treated groups suggestive of either interstitial or bronchopneumonia. Bronchoalveolar lavage (BAL) fluid, nasal and throat swabs were collected on Days 1, 3, and 6 after viral challenge (study Day 0). Viral genomes (gRNA) and subgenomic RNA (sgRNA), indicative of active viral replication (*12*), were detectable in BAL, throat swabs, and nasal swabs for all control animals following intranasal and intratracheal inoculation with SARS-CoV-2 (Fig. 4 and Fig. 5).

**Fig. 4.**
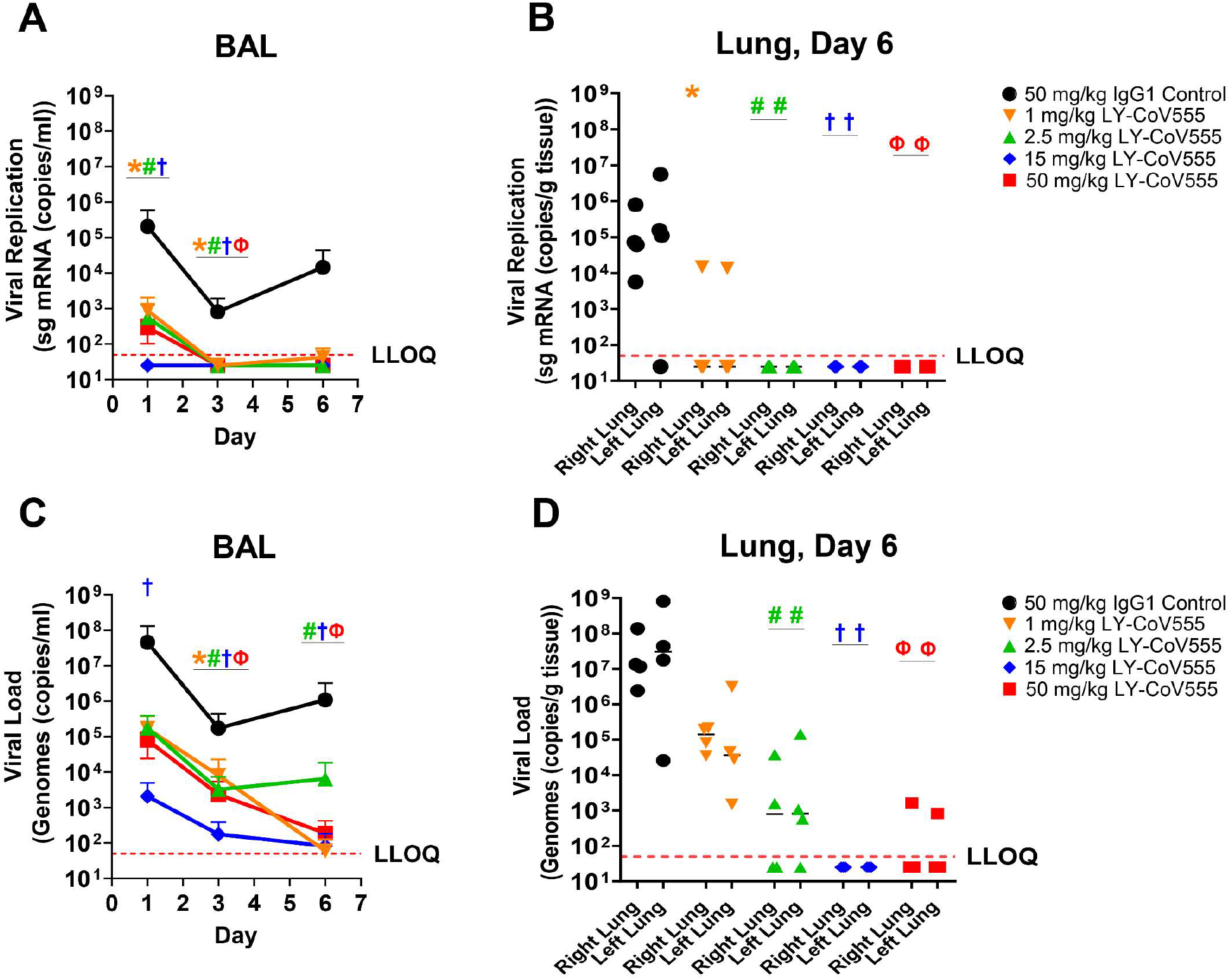
Impact of LY-CoV555 on lower respiratory tract viral replication and load in rhesus macaques challenged with SARS-CoV-2. 24 hours prior to viral challenge, rhesus macaques (N=3 or 4/group) were administered 1, 2.5, 15, or 50 mg/kg of LY-CoV555 as a single intravenous dose. sgRNA (viral replication) and gRNA (viral loads) were assessed by qRT-PCR in the BAL (A, C) over the course of 6 days post-inoculation. Viral replication and viral loads were assessed in lung tissue (B, D) h arvested on Day 6. A-C: Values represent the mean and standard error of the mean for 3 or 4 animals. D: bars represent the mean of 3 or 4 animals. Samples below the lower limit of quantification (LLOQ) were designated a value of ½ LLOQ. LLOQ = 50 copies for g or sg mRNA. Statistical analyses provided in Table S6. * denotes q-value <0.05, 1 mg/kg; ^#^ denotes q-value <0.05, 2.5 mg/kg; ^†^ denotes q-value <0.05, 15 mg/kg; ^Ф^ denotes q-value <0.05, 50 mg/kg. BAL = bronchoalveolar lavage; qRT-PCR = quantitative real-time polymerase chain reaction; sg mRNA = subgenomic messenger RNA.

**Fig. 5.**
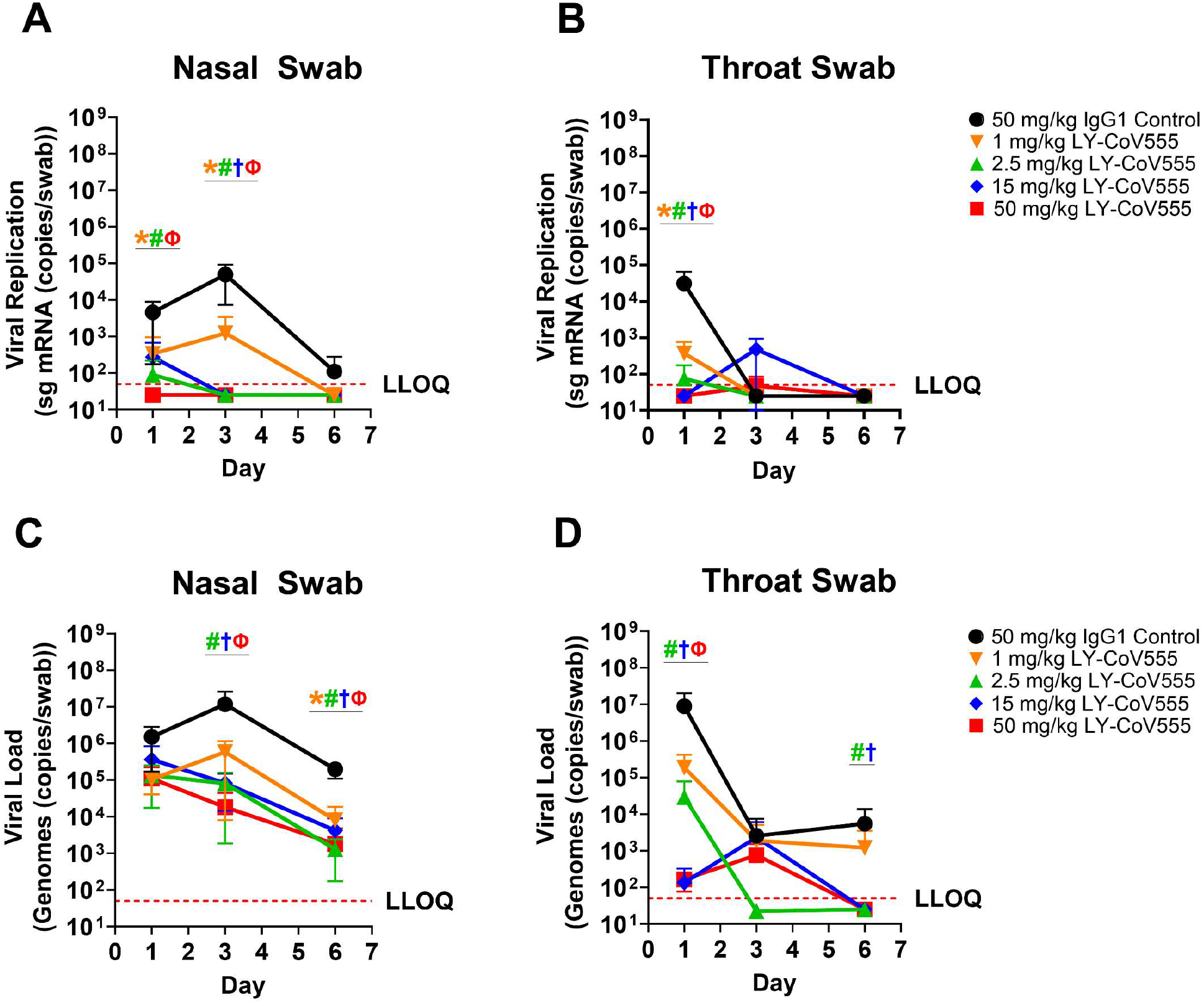
LY-CoV555 reduces viral replication and load in the upper respiratory tract. 24 hours prior to viral challenge, rhesus macaques (N=3 or 4/group) were administered 1, 2.5, 15, or 50 mg/kg of LY-CoV555 as a single intravenous dose. sgRNA (viral replication) and gRNA (viral loads) were assessed by qRT-PCR in the nasal swabs (A, C) and throat swabs (B, D) ove r the course of 6 days post-inoculation. A-C: Values represent the mean and standard error of the mean for 3 or 4 animals. D: bars represent the mean of 3 or 4 animals. Samples below the lower limit of quantification (LLOQ) were designated a value of ½ LLOQ. LLOQ = 50 copies for genomes or sg mRNA. Statistical analyses provided in Table S6. * denotes q-value <0.05, 1 mg/kg; ^#^ denotes q-value <0.05, 2.5 mg/kg; ^†^ denotes q-value <0.05, 15 mg/kg; ^Ф^ denotes q-value <0.05, 50 mg/kg. qRT-PCR = quantitative real-time polymerase chain reaction.

Prophylactic treatment with LY-CoV555 resulted in significant decreases in viral load (gRNA) and viral replication (sgRNA) in the lower respiratory tract following SARS-CoV2 inoculation, based on determinations in BAL and lung tissue (Fig. 4, Table S6). In the BAL, reductions of 10^2^ to 10^5^ in viral replication and load were observed compared to controls across Days 1, 3 and 6, with significant reductions in viral replication (1, 2.5, and 15 mg/kg doses; q<0.05) and load (15 mg/kg dose) on Day 1 and at all dose levels on Day 3 (Fig. 4A, C). In LY-CoV555-treated animals, viral replication in BAL was undetectable by Day 3 at all dose levels (Fig. 4A). Consistent with BAL on Day 6, no viral replication was observed in lung tissue harvested at necropsy in the 2.5, 15, and 50 mg/kg dose groups, demonstrating a significant reduction (q-value<0.05) compared to control (Fig. 4B, Table S6). Viral loads in the BAL and lung on Day 6 were significantly reduced (q-value<0.05) at the 2.5, 15, and 50 mg/kg doses (Fig. 4C, D, Table S6).

LY-CoV555 also provided protection in the upper respiratory tract, with a significant reduction in gRNA (q-value <0.05) at the 2.5, 15, and 50 mg/kg doses in the throat on Day 1 and the nose on Days 3 and 6 (Fig. 5C, D, Table S6). Most importantly, viral replication was significantly reduced in the nose (1, 2.5, 50 mg/kg doses) and throat (1, 2.5, 15, and 50 mg/kg doses) on Day 1 (q-value<0.05) and by Day 3, was undetectable in the nose (<50 copies/swab) at the 2.5, 15, and 50 mg/kg dose levels (Fig. 5A, B, Table S6).

Our results demonstrate that treatment with LY-CoV555 provides substantial upper respiratory tract protection and indicate the potential for reduced viral shedding and transmission following treatment with a neutralizing antibody. Given the robust nature and route of administration of the viral inoculum in this model, we hypothesize that modest doses of LY-CoV555 could provide substantial clinical efficacy. To this point, in a recent study of several prototype SARS-CoV-2 DNA vaccines, administration of a 10-fold lower inoculum following immunization still resulted in detectable levels of sgRNA in the nose of most immunized rhesus macaques.(*15*) Overall, we show dose-related reductions in gRNA and sgRNA in the upper and lower respiratory tracts with maximal protection observed at doses of 2.5 mg/kg and above.

LY-CoV555 administration resulted in sustained serum concentrations after IV dosing, consistent with expected pharmacokinetics for human IgG in a NHP model (Table S5). Mean serum concentrations of LY-CoV555 on the day of viral challenge were 15 ±3, 38 ±14, 276 ±37 and 679 ±101 μg/mL at the 1, 2.5, 15, and 50 mg/kg dose levels, respectively. The dose responsive concentrations of serum LY-CoV555 were consistent with the dose-related reductions in viral loads in the lungs, throat, and nasal passages. Importantly, serum concentrations of LY-CoV555 at doses of 2.5 mg/kg and higher were associated with maximal protection in this rhesus infection model. The delayed impact on viral loads in nasal swabs could reflect slower distribution of antibody into the nasal epithelial lining fluid versus the lung or throat.

This study describes the isolation and characterization of a potent anti-spike neutralizing antibody, LY-CoV555, derived from a convalescent COVID-19 patient. LY-CoV555 was found to have RBD binding and ACE2 blocking properties and displayed high affinity and potency due to its unique SARS-CoV-2 spike protein-binding properties. In both *in vitro* assays with full virus and an NHP model of SARS-CoV-2 infection, LY-CoV555 displayed high protection potency supporting its development as a therapeutic for the treatment and prevention of COVID-19. This study also provides evidence that neutralizing antibodies have potential as an important countermeasure to preventing and treating SARS-CoV-2 infection and reduce virus replication in the upper airway that may decrease transmission efficiency. LY-CoV555 is presently under clinical evaluation for the treatment and prevention of COVID-19 (NCT04411628; NCT04427501; NCT04497987; NCT04501978).

The importance of prototype pathogen preparedness was demonstrated by the ability to rapidly design and produce protein for B-cell probes based on prior work defining the structure and stabilization strategy for the betacoronavirus spike protein.(*24*) The resulting speed at which this drug discovery and development effort progressed, with achievement of first-in-human dose in only 90 days after the initiation of antibody screening, is a testament to the advanced discovery and characterization platforms and pre-established public-private partnership. Overall, the identification and characterization of LY-CoV555 points to the feasibility of strategies to rapidly identify neutralizing human mAbs as part of an initial response to an evolving pandemic that can complement population-scale vaccination, provide immediate passive immunity, and provide protection for vulnerable populations.

## Acknowledgments

We would like to thank the following: Kristi Huntington (Eli Lilly and Company) and Payal Sipahimalani (AbCellera Biologics Inc.) for project leadership and coordination; Douglas Burtrum, Nichole Niemela Mayer, Candyd Velasquez, Xiaomin Yang, Ricky Lieu, Richard Yuan, Dongmei He, Henry Koo, Michael Cohen, David Randolph, Paul Anderson, Regina White, Matt Schmitt, John Herrington, Rob Peery, Maria Hougland, Matt Jeffries, and Gavin Barnard of Eli Lilly and Company for reagent and antibody generation; Craig Dickinson, Kristina Coleman and Jeffrey Boyles of Eli Lilly and Company for initiation and reagent generation for crystallography experiments; Donald Lee and Anna Russell of Eli Lilly and Company for bioinformatic and structural analyses; Roy Heng and Jonathan Fitchett of Eli Lilly and Company for HDX data generation; Jenny Chien and Emmanuel Chigutsa of Eli Lilly for in vivo study design support; Catherine Brockus of Eli Lilly for bioanalytical support; Ross Blankenship and Gregory Dyas of Eli Lilly and Company for *in vivo* data analysis; Sherie Duncan, Anders Klaus, Keith Mewis, Karine Herve, Amanda Moreira, Aoise O’Neill, and Emilie Lameignere of AbCellera Biologics Inc. for technical support; Chad Thiessen of AbCellera Biologics Inc. for development of features for Celium™ required for antibody selection; Clara Ng-Cummings of AbCellera Biologics Inc. for figure generation; Hanne Andersen Elyard and Mark G. Lewis of BioQual Inc. for *in vivo* study conduct; Wolfgang Glaesner of Eli Lilly and Company for management/personnel resources; Suchetana Bhattacharyya and David McIlwain of Eli Lilly and Company for editorial and process support of this manuscript.

## Author contributions

B.E.J., P.B.A., C.M.W., J.D., T.P.C, J.L.B, B.A.H., D.B., D.F., M.H.P., J.H., R.E.H., and S.J.B. conceived of and designed experiments (LY-CoV555 *in vitro* and *in vivo* work), data analysis and reporting, and participated in manuscript authoring and review. F.J.T conceived of and designed experiments (mAb cloning) and participated in manuscript authoring and review. A.P. conceived of and designed experiments (crystallography/structure determination) and participated in manuscript authoring and review. A.C.A. participated in data analysis and reporting and manuscript review. J.R.M., B.S.G., K.S.C., L.W., and O.A. conceived of and designed experiments/reagents (pseudovirus neutralization assay), data analysis and reporting, and participated in manuscript authoring and review. R.S.B. and D.R.M. conceived of and designed experiments (nanoLuc assay), data analysis and reporting, and participated in manuscript review. M.J.M. conceived of and designed experiments, data analysis and reporting, and manuscript review. M.S. designed and prepared reagents for experiments and manuscript review. H.M.B. participated in data analysis and reporting and manuscript review. T.W.G., R.W.C., and V.B. conceived of and designed experiments (PRNT assay), data analysis and reporting, and participated in manuscript review. K.R.J. conceived of and designed experiments (ML-based mAb selection/ranking), data analysis and reporting, and participated in manuscript review. R.G. participated in conception and design of experiments (rapid mAb cloning, bioinformatics), data analysis and reporting. M.A.S. designed and implemented software improvements to Celium™ for antibody selection. R.G., M.A.S. and D.W.C., assisted in the acquisition, organization and interpretability of the data, and participated in manuscript review. S.J.H. and S.A.T. designed and developed features for Celium™ required for antibody selection and participated in manuscript review. M.D.V., A.D., and M.D. conceived of and designed experiments (I.F.A), data analysis and reporting, and participated in manuscript review. J.A.G., C.H., N.V.J., J.S.M., and D.W. conceived of and designed experiments (cryo-EM/mAb structure determinations), data analysis and reporting, and participated in manuscript review. S.Ž., K.W., and K.M. participated in design, execution, data analysis and interpretation (screening and validation experiments), as well as drafting and review of this manuscript. K.W. participated in the interpretation of screening, validation and characterization data for downselection of antibodies for expression and characterization. L.K. and Y.H. designed and executed binding kinetics & epitope binning experiments, data analysis and reporting, and participated in manuscript authoring and review. D.P. designed and implemented data analysis and reporting pipelines for binding kinetics, epitope binning and ACE2 blocking, and participated in manuscript authoring and review. R.B. conceived and designed experiments, analyzed and reported data (rapid mAb cloning) and participated in manuscript authorship. Y.H. participated in design, execution, and data analysis of rapid mAb cloning reported data. L.K. participated in design, execution and data analysis of mAb biophysical characterization data. P.X. conceived and designed experiments, generated the expression vectors for the full-length spike proteins (wildtype and mutants) for validation and participated in manuscript authorship. S.S. participated in data analysis and reporting (discovery and characterization) and manuscript authorship and review. B.C.B., C.L.H. and E.F. conceived and designed experiments, analyzed and reported data (discovery, characterization, bioinformatics and antibody lead selection) and participated in manuscript authorship and review.

## Disclaimer

This research was funded in part by the U.S. Government. The views and conclusions contained in this document are those of the authors and should not be interpreted as representing the official policies, either expressed or implied, of the U.S. Government. Approved for Public Release, Distribution Unlimited.

## Data and materials availability

All data associated with this study is available in the main text or the supplementary materials. Atomic coordinates and cryo-EM maps of the reported structure have been deposited in the Protein Data Bank under accession code XXX and in the Electron Microscopy Data Bank under accession code XXX.

## Supplementary Materials

Materials and Methods

Supplementary Text

Figs. S1 to S7

Tables S1 to S8

References (*25-56*)

