## Supplementary Materials for "LY-CoV555, a rapidly isolated potent neutralizing antibody, provides protection in a non-human primate model of SARS-CoV-2 infection"

**This PDF file includes:**

Materials and Methods

Supplementary Text

Figs. S1 to S7

Tables S1 to S8

References (*25-56*)

#### Materials and Methods

##### *Single-cell screening and recovery*

A blood sample from a 35-year-old male patient hospitalized (ICU) with COVID-19 was obtained mid-February 2020, approximately 20 days following the onset of symptoms. Peripheral blood mononuclear cells were isolated and cryopreserved using standard protocols. Cells were thawed, activated in culture to generate memory B-cells, and enriched for antibody-secreting B-cells prior to injection into AbCellera's microfluidic screening devices with either 91,000 or 153,000 individual nanoliter-volume reaction chambers. Single cells secreting target-specific antibodies were identified and isolated using two assay types (*25*) (Hansen, C. L. G., Lecault, V., Piret, J. M. & Singhal, A. System and method for microfluidic cell culture. United States patent US 10,421,936. Sep 24, 2019; Singhal, A., Hansen, C. L. G., Schrader, J. W., Haynes, C. A. & Costa, D. J. D. Methods for assaying cellular binding interactions. United States patent US 10,466,241. Nov 5, 2019): a multiplexed bead assay using multiple optically-encoded beads, each conjugated to the soluble pre-fusion stabilized spike of either SARS-CoV or SARS-CoV-2 spike with T4-foldon domain, 3C protease cleavage site, 6x His tags, and twin-strep tags (*24*) or negative controls (bovine serum albumin [BSA] His-tag and T4 FoldOn trimerization domain), and a live cell assay using passively dyed suspension-adapted Chinese hamster ovary (CHO) cells transiently transfected to surface-express full-length SARS-COV-2 spike protein (GenBank ID MN908947.3) with a green fluorescent protein (GFP) reporter, and non-transfected cells as a negative control. Beads or cells were flowed onto microfluidic screening devices and incubated with single antibody-secreting cells, and monoclonal antibody binding to cognate antigens was detected via a fluorescently labeled anti-human immunoglobulin G (IgG) secondary antibody. Positive hits were identified using machine vision and recovered using automated robotics-based protocols.

##### *Single-cell sequencing, bioinformatic analysis, and cloning*

Single cell polymerase chain reaction (PCR) and next-generation sequencing (MiSeq, Illumina) were performed using automated workstations (Bravo, Agilent) and custom molecular biology protocols for the recovery of paired heavy and light chain sequences. Sequencing data were analyzed using a custom bioinformatics pipeline to yield paired heavy and light chain sequences for each recovered antibody-secreting cell. Each sequence was annotated with the closest germline (V(D)J) genes, degree of somatic hypermutation, and potential sequence liabilities. Antibodies were considered members of the same clonal family if they shared the same heavy and light V and J genes and had the same CDR3 length. The variable (V(D)J) region of each antibody chain was PCR amplified and inserted into expression plasmids using a custom, automated high-throughput cloning pipeline. Plasmids were verified by Sanger sequencing to confirm the original sequence previously identified by next-generation sequencing.

##### *High-throughput antibody expression and purification*

Antibody-encoding plasmid DNA was transfected into Expi293-F™ cells (Thermo Fisher Scientific) using the manufacturer's recommended protocol with some modifications. Expi293-F™ cells were aliquoted into 24-well plates and incubated with shaking at 37°C with 8% $CO_2$ and 85% humidity. Heavy and light chain plasmids were pooled, mixed with Expi293Fectamine, and added dropwise to cells in each well. Expi293F Enhancer I and II were added 20 hours post-transfection and cell supernatants were harvested 4 days post-transfection. Antibody titers were measured by biolayer interferometry on an Octet HTX instrument (ForteBio).

Antibodies were purified using a standard protein A-based purification protocol and quantified by bio-layer interferometry using an Octet HTX instrument (ForteBio). The percent purity and apparent molecular weight of antibodies were analyzed by denaturing capillary electrophoresis SDS page using a LabChip GXII Touch instrument (Perkin Elmer) according to the manufacturer's protocol.

Larger scale antibody production utilized expression vectors for both chains contained inverted repeat sequences needed for chromosomal integration with the PiggyBac Transposase (Transposagen). DNA fragments corresponding to the variable regions of the antibodies of interest were either synthesized as eBlocks (IDT DNA), or were amplified from bulk plasmid DNA preparations using Phusion DNA Polymerase (NEB), with primers hybridizing to the FR1 and J-regions of the antibody with the same vector flanking sequences described above. Variable domain fragments were incorporated into the full length respective heavy and light chain expression vectors by Gibson Assembly (*26*) and transformed into *E. coli* (DH5a) for plasmid propagation and sequence confirmation. Sequence-confirmed plasmids containing the variable light chain and its cognate variable heavy chain (IgG1 constant domain) were mixed together into 96-deep well Master-blocks (Greiner) using the Biomek i7 Automated Liquid Handling Workstation (Beckman Coulter) for transfection into CHO cells.

CHO cells were maintained and transiently transfected as described previously (*27*), with the following modifications. Parental CHO cells were transfected in CD-Forti CHO (ThermoFisher) supplemented with 8mM L-Glutamine (Sigma-Aldrich) and 0.125% *N*,*N* dimethylacetamide (Sigma-Aldrich) at a final density of $4 \times 10^6$ viable cells/mL. Plasmids encoding both chains of each antibody were co-transfected with a plasmid encoding PiggyBac Transposase (Transposagen). Bulk stable cell pools expressing each antibody were generated by transferring 200µL from the transient CHO cultures into a new 24DWP containing 2 mL per well of selection LM (no L-glutamine, 25µM Methionine Sulfoximine [MSX]). Cells were recovered and fed batched as described in previously.(*28*)

Conditioned media supernatants containing the expressed antibodies were harvested by centrifugation, and expression titers were measured in the conditioned media using Protein G (ProG) Dip and Read Biosensors (FortéBIO) on a high throughput Octet HTX system (FortéBIO). Expressed antibodies were purified using mAb Select (Protein A) resin (Amersham Bio) in 24-deep well plate format using automated liquid handling and vacuum filtration. Purified antibodies were transferred to a 96-well round bottom plate (Costar), and concentrations were determined by 280 nm absorbance using a DropSense96 (Perkin Elmer), followed by normalization to 1 mg/mL using a Lynx LM I800 VVP (Dynamic Devices). Subsequent larger scale production of antibodies utilized larger volume transient transfections or larger volume production from stable cell lines, followed by purification using standard approaches. Fab portions of antibodies were generated by proteolytic digestion using Papain, followed by removal of un-cleaved protein using standard chromatography techniques.

The pre-fusion stabilized form of the SARS-CoV-2 spike protein ectodomain was used for binding studies.(*24*) The protein was produced in mammalian cells by transient transfection and purified; recombinant protein was obtained either produced from HEK293cells or CHO cells. The isolated receptor-binding domain (RBD) (using residues 328-541), was fused to a linker sequence containing a TEV-protease recognition site, followed by a human IgG1 Fc sequence. The monomeric spike RBD was generated by TEV protease digestion of the corresponding Fc-fusion reagents (transiently expressed in CHO cells as above, and purified by standard techniques), followed by purification of the isolated domains away from free Fc and un-cleaved Fc-fusion. The His-tagged angiotensin converting enzyme 2 (ACE2) ECD was expressed similarly in CHO cells, and purified by standard approaches.

*Binding validation and analysis*

Recombinant antibodies were confirmed to bind screening targets using two assay types via high-throughput flow cytometry. In a multiplexed bead-based assay, optically encoded beads were conjugated to one of the following unique antigens: spike proteins of SARS-CoV-2, MERS-CoV, SARS-CoV, HKU1-CoV, WIV1, or the S1 subunit of SARS-CoV-2 spike protein. Purified antibodies were incubated with target-conjugated and negative control BSA His-tag and T4 FoldOn-conjugated beads at either 50 nM, 10 nM or 2 nM antibody concentration for 30 minutes at room temperature. In a live cell-based assay, full-length spike protein sequences of either the wildtype or mutants V367F, V483A, and D614G of SARS-CoV-2 with GFP inserts were transiently transfected into CHO cells (MaxCyte STX Scalable Transfection System). Full-length native conformation spike protein expression was confirmed via GFP detection, flow cytometry-detected binding to S1 and S2 subunit-specific benchmark antibodies, and by western blot. Purified antibodies were incubated with target-expressing cells and non-transfected control cells at 50 nM, 10 nM, or 2 nM antibody concentration for 30 minutes at 4°C. Beads or cells were washed, and binding was detected using a fluorescently labeled anti-human IgG secondary antibody. Fluorescence was measured using high-throughput plate-based flow cytometry. Benchmark antibodies previously identified from a SARS-CoV convalescent patient sample and cross-reactive to SARS-CoV-2 spike protein were used as positive controls; human IgG isotype and an irrelevant antibody (HIV VRC01) were used as negative controls. Median fluorescence intensity of each antibody was normalized over the median fluorescence intensity of the human isotype, with signals greater than 5-fold over isotype control (and less than 2.5-fold binding to negative controls) considered as specific binding.

*Differential scanning fluorimetry*

Melting temperatures ($T_M$) of antibodies (175 µg/mL) was assessed by differential scanning fluorimetry using the SYPRO™ Orange fluorescence probe (10-fold molar excess, Thermo Fisher Scientific). Thermal unfolding as assessed by a change in fluorescence was measured on a Bio-Rad C1000 Touch

Thermal Cycler instrument (Bio-Rad Laboratories) using a CFX96 Real-Time System reader head (Bio-Rad Laboratories). The wavelengths for excitation and emission were 450 to 490 nm and 560 to 580 nm, respectively. Fluorescence signal was measured from 25 to 95 °C and melting curves were integrated using the Bio-Rad CFX Maestro software (v 1.1). The $T_M$ was defined as the local minimum obtained from the derivative of the melting curve.

*Dynamic light scattering measurements*

Percent aggregation and polydispersity of antibodies were assessed by dynamic light scattering (on a DynaPro® Plate Reader III instrument (Wyatt Technology) at varying antibody concentrations (0.5 – 2 mg/mL). DLS was carried out at 20 °C with 5 x 5 seconds acquisitions per individual sample. Data were analyzed in the Dynamics software (Wyatt Technology, v 7.9.0.5) using the regularization algorithm. Percent polydispersity and percent mass of soluble antibodies were calculated for the size range of 2 to 10 nm.

*Surface-plasmon resonance binding experiments*

All high-throughput surface plasmon resonance (SPR) binding, epitope binning and ACE2 competition experiments were performed on a Carterra® LSA™ instrument equipped with an HC-30M chip type (Carterra-bio) using a 384-ligand array format. For all experiments, antibodies were coupled to the HC-30M chip: the chip surface was first activated by flowing a freshly prepared 1:1:1 activation mix of 100 mM MES pH 5.5, 100 mM S-NHS, and 400 mM EDC for 7 minutes, and antibodies diluted to either 10 µg/mL or 1 µg/mL in 10 mM NaOAc pH 4.25 buffer 0.01% Tween were injected and printed simultaneously onto the chip surface for 10 minutes by direct coupling. The chip surface was quenched by flowing 1 M EtOHamine for 7 minutes, followed by 2 wash steps of 15 seconds each in 25 mM MES pH 5.5 buffer. Relevant benchmarks and negative control antibodies (HIV VRC01, mouse FoldOn 8203-C1, and rabbit His-tag PA1-983) were also printed on the chip surface.

For binding kinetics and affinity measurements, a 3-fold dilution series of the antigen of interest, starting at 300 nM in HBSTE + 0.1% BSA running buffer was sequentially injected onto the chip surface. For each concentration, the antigen was injected for 5 minutes (association phase), followed by running buffer injection for 15 min (dissociation phase). Two regeneration cycles of 15 seconds were performed between each dilution series by injecting Pierce IgG elution buffer (Thermo Fisher) + 1 M NaCl on the chip surface. The data were analyzed using the Carterra Kinetics analysis software using a 1:1 Langmuir binding model to determine apparent association ($k_a$) and dissociation ($k_d$) kinetic rate constants and binding affinity constants ($K_D$).

For epitope binning experiments, antibodies coupled to the chip surface were exposed to various antibody:antigen complexes. Samples were prepared by mixing each antibody in 10 to 20-fold molar excess with antigen (1:1 freshly prepared mix of 400 nM antibody and 40 nM antigen, both diluted in 1X HBSTE + 0.1% BSA running buffer). Each antigen-antibody premix was injected sequentially over the chip surface for 4 minutes (association phase to ligand printed onto chip previously), followed by a running buffer injection for 2 minutes (dissociation phase). Two regeneration cycles of 15 seconds were performed between each premix sample by injecting 10 mM glycine pH 2.0 onto the chip surface. An antigen-only injection (20 nM concentration in running buffer) was performed every 8 cycles. The data were analyzed using the Carterra Epitope analysis software (version 1.2.0.1960) for heat map and competition network generation. Analyte binding signals were normalized to the antigen-only binding signal, such that the antigen-only signal average is equivalent to one RU (relative unit). A threshold window ranging from 0.9 RU to 1.1 RU was used to classify analytes into 3 categories: blockers (binding signal under the lower limit threshold), sandwichers (binding signal over the higher limit threshold) and ambiguous (binding signal between limit thresholds). Antibodies with low coupling to the chip, poor regeneration or with absence of self-blocking were excluded from the binning analysis. Like-behaved antibodies were automatically clustered to form a heat map and competition plot.

For ACE2 competition, antibodies coupled to the chip were exposed to spike protein:ACE2 complex; 20nM of SARS-CoV-2 spike protein was premixed with 200 nM of the His-tagged ACE2 (ACE2-His) diluted in HBSEP+ with 0.5 M NaCl, 1% BSA, 1x Dextran, and 2 mg/mL Heparin, and incubated for about 12 hours. The complex of spike protein/ACE2-His was then tested for binding to immobilized antibodies on the prepared HC30M chip, with association for 5 minutes and dissociation for 1 minute. Regeneration was performed in 20mM Glycine pH 2.0 with 1 M NaCl for 30 seconds twice.

*Biolayer interferometry binding experiments with Fabs*

Biolayer Interferometery (BLI) experiments used the Fab portion of antibodies (Fig. S6). 50 nM 2X-Strep-tagged SARS-CoV-2 ectodomain in BLI buffer (10 mM HEPES pH 7.5, 150 mM NaCl, 3 mM EDTA, 0.05% Tween 20 and 1 mg/mL BSA) was immobilized onto a Streptavidin biosensor (FortéBio) for 600s using an Octet RED96e (FortéBio). The biosensor was then dipped into 100 nM Fab (diluted in BLI buffer), and the association signal was measured for 600 seconds. Following this, the biosensor was dipped into BLI buffer to measure the dissociation signal for 600 seconds. Data were reference-subtracted and fit to a 1:1 binding model using Octet Data Analysis Software v11.1 (FortéBio).

*Hydrogen-deuterium exchange followed by mass spectrometry*

Hydrogen-deuterium exchange mass spectrometry was performed in order to determine structural location of the binding site for antibodies on the SARS-CoV-2 spike protein. Instrumentation used for the experiments is as described by Espada et. al. (*29*) with the sample injected directly into a LEAP PAL3 HDX autosampler and resolved using a Waters Synapt G2Si (Waters Corporation). Peptide identification used 5 μg spike protein (1:10 dilution in 0.1 X phosphate buffered saline in $H_2O$) using nepenthesin II for digestion in HDMSe (Mobility ESI+ mode). Complexes of human SARS-CoV-2 spike protein with individual antibodies was prepared at the molar ratio of 1:1.2 in 10 mM sodium phosphate buffer, pH 7.4 containing 150 mM NaCl (1xPBS buffer). The exchange experiment was initiated by adding 25μL of $D_2O$ buffer containing 0.1x PBS to 2.5 μL of spike protein (1 mg/ mL) or spike- antibody complex at 15˚C for various amounts of time (0 seconds, 10 seconds, 30 seconds, 2 minutes, 10 minutes, and 120 minutes). The peptides derived from samples of the free and bound states of SARS-CoV-2 spike protein were compared for deuterium incorporation differences to identify protected regions indicative of the binding epitope as described.(*30*)

*Negative-stain electron microscopy*

SARS-CoV-2 spike ectodomain was diluted to 0.04 mg/mL in 2 mM Tris pH 8.0, 200 mM NaCl, 0.02% $NaN_3$ (dilution buffer) in the presence of 10-fold excess Fab and incubated on ice for 10 seconds. CF400-Cu grids (Electron Microscopy Sciences) were plasma cleaned for 30 seconds. in a Solarus 950 plasma cleaner (Gatan) with a 4:1 ratio of $O_2/H_2$. A volume of 4.8 μL of the protein sample was applied to the grid and allowed to incubate for 30 seconds. The grid was then washed twice with dilution buffer prior to staining with methylamine tungstate (NANO-W, Nanoprobes). Grids were imaged using a FEI Talos TEM (Thermo Scientific) and a Ceta 16M detector. Micrographs were collected manually using TIA v4.14 software at a magnification of ×92,000, corresponding to a pixel size of 1.63 Å/pixel. Contrast transfer function (CTF) estimation and particle picking were performed in cisTEM. A 2D classification was performed in either cisTEM (*31*) or cryoSPARC v2.15.10 (*32*), and antibody initio reconstruction and refinement of 3D maps were performed in cryoSPARC.

*Cryo-electron microscopy*

A purified, prefusion stabilized SARS-CoV-2 spike variant, HexaPro(*33*) at 0.2 mg/mL was complexed with 1.3-fold molar excess of LY-CoV555 Fab in 2 mM Tris pH 8, 200 mM NaCl, 0.02% $NaN_3$ for 5 minutes on ice. Three microliters of protein complex were deposited on an UltrAuFoil 1.2/1.3 grid (Electron Microscopy Sciences) which had been plasma cleaned for 2 minutes using a Gatan Solarus 950 with a 4:1 $O_2$:$H_2$ ratio. The grid was then plunge-frozen in liquid ethane using a Vitrobot Mark IV (Thermo Scientific) set to 100% humidity and 22 °C, with a blot time of 5 seconds and a blot force of -4. Data were collected on a Titan Krios operating at 300kV and equipped with a K3 detector using a magnification of 22,500x, resulting in a pixel size of 1.045 Å. A total of 30 frames were collected for each micrograph, with defocus values ranging from -0.8 µm to -2.8 µm, a total exposure time of 4.5 seconds, and a total electron dose of ~32.7 $e^-/Å^2$. A full description of the data collection parameters can be found in Table S7 and Figure S7. Motion correction, CTF estimation, and particle picking were performed in Warp.(*34*) Particles were subsequently transferred to cryoSPARC v2.15.10 (*32*) for 2D classification and 3D reconstruction. The refined map was then subjected to local B-factor sharpening using LocalDeBlur.(*35*) Model building and refinement were subsequently performed using Coot, Phenix and ISOLDE.(*36-38*)

##### *Protein crystallography*

For protein crystallography, an isolated RBD (using residues 329 to 527), was fused to a 6-His tag at the C-terminus, expressed in CHO cells, enzymatically deglycosylated using endoglycosidase-H, and purified by cation exchange chromatography. The Fab portions of selected antibodies, containing mutations in the constant region known to encourage crystallization (*39*), were expressed in CHO cells, and purified. The Fab:RBD complexes were prepared by mixing the components, with a 20% excess of the RBD, and then the complex purified from the excess RBD by size-exclusion chromatography. Fab:RBD complexes (approximately 12 mg/mL) were crystallized by vapor diffusion sitting drops. Crystals of complexes formed within 1 to 2 days and were harvested on the third day. Crystals were flash-frozen in liquid nitrogen following 1-minute incubation in cryoprotectant solution containing 25 % glycerol in mother liquor: LY-CoV555 Fab-RBD complex crystallized using 100 mM sodium acetate pH 4.6 and 20 % PEG 10K; the 481CK Fab-RBD complex crystallized using 100 mM Tri-Sodium Citrate pH=5.8, and 14% PEG 4K, and 10% 2-Propanol; and the 488 CK Fab-RBD complex crystallized using 100 mM Hepes pH=7.7, and 8% PEG 3350, and 200 mM L-Proline.

Diffraction data were collected at Lilly Research Laboratories Collaborative Access Team and beamline at Sector 31 of the Advanced Photon Source at Argonne National Laboratory, Chicago, Illinois. Crystals stored in liquid nitrogen were mounted on a goniometer equipped with an Oxford Cryosystems cryostream maintained at a temperature of 100 K. The wavelength used was 0.9793 Å, collecting 900 diffraction images at a 0.2-degree oscillation angle and 0.12 seconds exposure time on a Pilatus3 S 6M detector at a distance of 392 mm. The diffraction data were indexed and integrated using autoPROC (*40*) / XDS (*41*) and merged and scaled in AIMLESS (*42*) from the CCP4 suite.(*43*) Non-isomorphous data readily yielded initial structures by Molecular Replacement using for the Fab portion crystal structures from the proprietary Eli Lilly structure database and for the SARS2 spike RBD from the public domain structure with the access code 6yla (Huo et al. 2020; manuscript in preparation). The initial structure coordinates for each dataset were further refined using Refmac5 (CCP4) applying isotropic temperature factors. Model building was performed with Coot (CCP4) and final structure validation with MolProbity (*44*) and CCP4 validation tools. Table **S8** presents the crystallographic data statistics.

Protein coordinates and structure factors have been deposited with the Protein Data Bank under the access codes 1xxx, 2xxx, 3xxx.

*Pseudotyped neutralization assay for monoclonal antibody screen*

SARS-CoV-2 spike pseudotyped lentiviruses that harbor a luciferase reporter gene were produced and neutralization assay was performed as described previously.(*45, 46*) Pseudovirus was produced by co-transfection of 293T cells with plasmids encoding the lentiviral packaging and luciferase reporter, a human transmembrane protease serine 2 (TMPRSS2), and SARS-CoV-2 S (Wuhan-1, Genbank #: MN908947.3) genes. Forty-eight hours after transfection, supernatants were harvested, filtered and frozen. For initial screening neutralization assay 4 dilutions of monoclonal antibodies (10, 1, 0.1, and 0.01ug/mL) were mixed with titrated pseudovirus, incubated for 45 minutes at 37 °C and added to pre-seeded ACE2-transfected 293T cells (either transiently or stably transfected) in 96-well white/black Isoplates (Perkin Elmer). Following 2 hours of incubation, wells were replenished with 150 µL of fresh medium. Cells were lysed 72 hours later and luciferase activity (relative light unit, RLU) was measured. Percent neutralization was calculated relative to pseudovirus-only wells.

*Neutralization of virus activity*

Viral neutralization activity of the discovered antibodies was measured by detecting the neutralization of infectious virus in cultured Vero E6 cells (African Green Monkey Kidney; ATCC #CRL-1586). These cells are known to be highly susceptible to infection by SARS-CoV-2. Cells were maintained according to standard ATCC protocols. Briefly, Vero E6 cells were grown in Minimal Essential Medium (MEM) supplemented with 10% heat-inactivated fetal bovine serum (FBS), 2mM L-glutamine, and 1% of MEM Nonessential Amino Acid (NEAA) Solution (Fisher #MT25025CI). Cell cultures were grown in 75 or 150 $cm^2$ flasks at 37°C with 5% $CO_2$ and passaged 2 to 3 times per week using trypsin-EDTA. Cell cultures used for virus testing were prepared as subconfluent monolayers. All incubations containing cells were performed at 37°C with 5% $CO_2$.

*Production of virus Inocula*

Immunofluorescent and plaque reduction assays were conducted using virus produced by infecting cultured Vero E6 cells with the SARS-CoV-2 clinical isolate USA/WA/1/2020 (BEI resources number NR52281) or the Italy-INMI1 isolate (European Virus Archive – Global, ref #008V-03893) and incubating at 37ºC until cytopathology is evident (typically 48 to 72 hours). Expansion was limited to 1 to 2 passages in cell culture to retain integrity of the original viral sequence. The virus stock was quantified by standard plaque assay, and aliquots were stored at -80ºC. A freshly thawed aliquot was used for each neutralization experiment.

*Virus neutralization detected by immunofluorescence*

Virus infectivity assays were conducted in 96-well tissue culture plates. Vero E6 cells were seeded at a density of $8x10^4$ cells/cm$^2$ and incubated overnight to a confluency of approximately 95%. Serial dilutions of antibodies or positive control polyclonal serum from a convalescent SARS-CoV-2 patient, were prepared in DMEM (Dulbecco's Modified Essential Medium, Gibco # 11965-092) supplemented with 1% NEAA and 10mM HEPES. Virus stock (prepared for a final concentration of 18 to 20 TCID50 per well) was added to each dilution of antibody and incubated for 1 hour. Virus with no antibody and a no-virus wells served as controls. Incubated samples were inoculated onto Vero E6 cell at a final volume of 100ul, and plates were incubated 24 hours. To detect virus replication, the inoculum was removed, and monolayers were fixed in 10% formalin solution (4% active formaldehyde) for 1 hour at room temperature (RT). Background staining was quenched by adding 50 mM $NH_4Cl$ to cells and rocking for 10 minutes at RT, followed by washing. Cells were permeabilized with 0.1% Triton-X 100 (by rocking at RT for 10 min), washed 3X with DPBS, and nonspecific antibody binding was blocked with 1% BSA. Mouse anti-SARS-CoV-2 nucleoprotein antibody (1 C7C7, Dr. Thomas Moran, Icahn Sch of Med, Mount Sinai), diluted at 1:1000 in DPBS with 1% BSA, was added to each well and incubated overnight at 4°C. After washing, cells were stained with goat anti-mouse Alexa Fluor plus 647 antibody (Thermo Fisher #

A32728; green dye) and DAPI (4',6-diamidino-2-phenylindole, dihydrochloride; Thermo Fisher # 62247; blue dye) by incubating for 1 hour at 37°C. Images were collected using a CellInsight CX7 with the 4× objective covering the entire well. The percentage of infected cells per well relative to the uninfected and no-antibody controls was analyzed using the instrument's "Target Activation" analysis protocol.

*Virus neutralization detected by luciferase reporter*

Luciferase assays were performed using a molecular complementary DNA clone of a SARS-CoV-2 isolate (USA/WA/1/2020) in which a non-essential gene (ORF7) was replaced by the NanoLuc luciferase reporter gene (Promega), as previously described for SARS-CoV and MERS-CoV (Sheahan et al, and references therein).(*47*) Virus infectivity assays were conducted in 96-well tissue culture plates. Vero E6 cells were seeded at a density of $2x10^4$ cells per well in DMEM medium supplemented with 10% FBS (DMEM/FBS) and incubated for 15 to 24 hours. The next day, serial dilutions of antibodies or hIgG1 isotype control were prepared in DMEM/FBS. The SARS-CoV-2-NanoLuc inoculum was diluted in DMEM/FBS, mixed with an equal volume of diluted antibody (to produce a final virus titer of 140 plaque-forming unit [pfu]/well), and incubated 1 hr. After removing the culture medium from the plated Vero E6 cells, the virus-antibody solution was inoculated onto duplicate wells and incubated for 48 hours. Following standard protocols as recommended by the vendor, NanoGlo reagent (Promega #N1110) was added and luciferase activity was quantified on a SpectraMax plate reader (Molecular Devices).

*Virus neutralization detected by plaque reduction*

Plaque reduction assays were performed in 6-well plates. Vero E6 cells were seeded at a concentration of approximately $10^6$ cells/well and grown overnight to reach 95% confluency. The next day, serial three-fold dilutions of antibody were prepared in Eagle's MEM, mixed with approximately 100 pfu of SARS-CoV-2, and incubated for 1 to 2 hours. The antibody/virus mixtures were inoculated directly onto the cells and allowed to adsorb for 1 hour, with rocking at 15-minute intervals. An overlay media composed of 1.25% Avicel RC-581 (FMC BioPolymer) in Eagle's MEM with 5% FBS was added, and plates were incubated for 48 (Italy isolate) or 72 hours (USA/WA1 isolate) for virus plaques to develop. After incubation, overlays were removed by aspiration and the cells were fixed with 10% buffered formalin-containing crystal violet stain for 1 hour. Plaques were counted manually, and plaque forming units were determined by averaging technical replicates per sample. Percent neutralization was determined relative to IgG isotype antibody control-treated wells.

*Non-human primate challenge*

The rhesus macaque model of SARS-CoV-2 infection was conducted according to the method of Chandrashekar et al.(*12*). This study was approved by the Institutional Animal Care and Use Committee of BioQual Inc. in accordance with the animal welfare requirements and accreditations. Housing and handling of the animals was performed in accordance with the standards of the AAALAC International's reference resource: the eighth edition of the *Guide for the Care and Use of Laboratory animals*, Animal Welfare Act as amended, and the 2015 reprint of the Public Health Service (PHS) Policy on Human Care and Use of Laboratory Animals. Handling of samples and animals was compliance with the Biosafety in Microbiological and Biomedical Laboratories (BMBL), 5$^{th}$edition (Centers for Disease Control). Naïve female rhesus macaques of Indian origin (purpose bred, Macaca mulatta from PrimGen 8 to 12 years of age) were administered at 1, 2.5, 15, or 50 mg/kg of LY-CoV555 or 50 mg/kg of an IgG1 control antibody by slow intravenous bolus (N= 3 or 4 animals per group). On study Day 0 (one day following antibody administration), monkeys received a viral challenge of 1.1 x$10^5$ PFU SARS-CoV-2 USA-WA1/2020 in 2 mL volume administered divided as 0.5 mL/nostril (IN) and 1.0 mL intratracheally (IT). Live phase parameters were monitored pre-study through necropsy (Day 6). COVID specific observations were collected daily in conscious animals to monitor overall health and welfare and determine the need for veterinary intervention and/or euthanasia. COVID observations were scored on a scale of 0 to 10 and included measures of respiratory rate and dyspnea, overall appearance, activity, and responsiveness. Clinical observations were assessed cage side twice daily and included evaluations of overall animal appearance, fecal consistency, and appetence. Body weights and rectal body temperatures were measured daily in anesthetized animals. At termination on Study Day 6, macroscopic observations in the lung were evaluated by a board-certified veterinary pathologist.

Bronchioalveolar lavage (BAL), nasal and oral swabs were collected on Days 1, 3 and 6, and lung tissue samples (at necropsy, Day 6) were collected to assess genomic and subgenomic viral RNA via qRT-PCR, conducted as reported(*12, 15*). The lower limit of detection for genomic and sub-genomic RNA copies was 50. In cases where the values were below the lower limit of detection in the assay, a value of 25 (1/2 the limit of quantitation) was used for calculations. Serum samples were also collected for determination of LYCoV-555 concentrations by total human IgG ELISA assay.

*Serum ELISAs for human IgG concentrations*

Concentrations of human IgG in rhesus macaque serum were determined by an ELISA assay. Goat anti-Human Kappa Monkey ads-UNLB (Southern Biotech, Catalog Number 2064-01; 1.00 µg/mL) was coated on the ELISA plate (ThermoFisher Scientific, Catalog Number 3855 or equivalent) as the capture reagent. Calibrators, controls and samples in neat rhesus macaque serum were diluted 200-fold, were transferred to the coated plates. After incubation, the plate was washed to remove unbound material, and mouse anti-human IgG Fc -HRP (Southern Biotech, Catalog Number 9040-05; 10 ng/mL) was added as detection reagent. Following incubation, unbound enzyme was washed away and BioFX® TMB One Component HRP Microwell Substrate (SurModics, Catalog Number TMBW-0100-01 or equivalent) was added to the wells. Color development was stopped by the addition of Phosphoric Acid (Fisher Chemical,

Catalog Number A260-500 or equivalent) and the optical density was measured at 450 nm with wavelength correction set to 650 nm. Immunoreactivity was determined from calibrators using a 4-parameter logistic (Marquardt) regression model with 1/F2 weighting (Watson Bioanalytical LIMS, version 7.4.2 SP1).

*Statistical analysis*

*In vitro* neutralization potencies were estimated using percent neutralization, log10 transformed antibody concentration, and a four-parameter logistic model fit using the drc() package (*48*) with R version 3.6.3.(*49*) All four parameters were estimated from the fitting and neutralizing concentrations were reported using absolute neutralization levels. Overall potency estimates were obtained by meta-analysis of all SARS-CoV-2 neutralization potency estimates using a random effects model with the metafor R package.(*50*)

Due to the left-censored nature of the rhesus macaque viral load data, study sample size, and the need for multiple comparisons correction due to the number of tests being conducted, a multiple imputation approach was favored over a non-parametric testing strategy. Multiple imputation (m=20 imputations) was conducted in accordance with standard procedures described by Rubin.(*51*) All statistical analyses were done using log10 transformed viral load values as the response. Imputation of left-censored data was done using random normal values with variance matched to the non-censored viral load values. Following imputation, a standard MMRM (mixed model repeated measures) model was fit with lme (*52*) using animal as a random effect, group, day, and group*day as fixed effects, and an unstructured covariance matrix. Treatment effects were pooled in accordance with Rubin to estimate a pooled effect size, standard error, and p-values.(*51*) Pooled p-values were estimated from a t-distribution with the degrees of freedom derived from the method described by Barnard and Rubin.(*53*) Within study pooled p-values were then adjusted for multiplicity using the Benjamini Hochberg method.(*54*)

#### Supplementary Text

##### *Screening, Binding validation and characterization of recombinant antibodies*

We deployed two screening assays: (1) a multiplexed bead-based assay using optically-encoded microbeads, each conjugated to either soluble prefusion-stabilized trimeric SARS-CoV-2 or SARS-CoV spike protein to screen for both SARS-CoV-2 monospecific and SARS-CoV-2/SARS-CoV cross-reactive antibodies and (2) a live cell-based assay using mammalian cells that transiently express full-length membrane-anchored SARS-CoV-2 spike protein, expected to present epitopes of the trimeric spike protein that mimic the native virion-displayed conformations during *in vivo* infection (Fig. S1).(*55*) Next-generation sequencing (NGS) libraries of antibody genes from recovered single B cells were generated and sequenced, and a custom bioinformatics pipeline with machine learning (ML)-based sequence curation was used to recover paired-chain antibody sequences, resulting in 440 unique high-confidence heavy and light paired-chain sequences (Fig. 1, Fig. S1B).

From the 175 cloned antibodies, all were successfully expressed in mammalian cells with 161 having sufficient expression levels for biophysical and functional characterization (summarized in Fig. S2). The SARS-CoV-2 spike protein binding properties of the recombinant antibodies and cross-reactivity to other coronaviruses soluble spike proteins were validated using a multiplexed bead assay via automated high-throughput flow cytometry. Of the 175 selected antibodies, 92% of antibodies validated as SARS-CoV-2, 34% as bat SARS-like coronavirus WIV1, 31% as SARS-CoV, 3% as Human coronavirus HKU1, 2% Middle Eastern respiratory syndrome coronavirus (MERS-CoV), and 2% as cross-binders to all spike proteins (Fig. S2B). Furthermore, 51% of antibodies validated as SARS-CoV-2 S1 subunit-specific binders, with 8% cross-binding to full length WIV1 and 6% cross-binding to full-length SARS-CoV, suggesting that as expected, most cross-binders are S2 subunit-specific. Antibody binding to cell-expressed, full-length SARS-CoV-2 wild-type spike and known circulating variants (V367F, V483A, D614G) was validated via automated high-throughput flow cytometry (Figure S2B). In this assay format, 77% of antibodies were validated as wild-type binders. Of that subgroup, 93% also validated for binding to two RBD mutations (V378F and V483A) or the very common D614G non-RBD mutation. In addition, 76% of antibodies were validated in both multiplexed bead-based and live cell-based assays (Fig. S2B) indicating the robustness of the single-cell screening assays with integrated ML-based hit-detection for identifying SARS-CoV-2-specific antibodies. Consistent with the bead and cell-based binding studies, these antibodies exhibited high affinity binding to the soluble spike protein in surface plasmon resonance (SPR) capture kinetic experiments using a Carterra® LSA™ instrument (Fig. 2A, Fig. S2C). Of these, 53% of antibodies had apparent binding affinity constant ($K_D$) values in the picomolar range and the remaining 47% in the nanomolar range, with a mean $K_D$ value of 5.3 nM. Due to the trimeric nature of the soluble spike protein and the potential bi-valent binding by the coupled antibodies, these affinities are substantially greater than true monomeric binding affinities (Table S2), but likely are more representative of the pharmacological setting.

Epitope binning SPR experiments were used to characterize the epitope coverage of the discovered antibodies. Benchmark antibodies with known binding to S1, NTD, RBD, and S2 epitopes of the SARS-CoV spike protein and cross-reactivity to SARS-CoV-2 spike protein were included to mark epitope identity. Cross-blocking results are summarized in the competition plot (Fig. 2B), as well as in the heat map (Fig. S3). In total, 95 unique bins (including the controls) were identified and a clear divide between S1 and S2-specific antibodies as inferred by benchmark competition is seen (Fig. 2B), suggesting these antibodies possessed a broad epitope diversity. Consistent with this, we observed that only approximately 10% of the antibodies tested exhibited ACE2 competition. The recombinantly expressed antibodies were subsequently screened in a high throughput pseudotyped lentivirus reporter neutralization assay. A summary of the distribution of the observed neutralization activities from these antibodies (Fig. 2C) demonstrates that greater inhibition is linked to the ability to block interaction with ACE2 as expected, but that antibodies to other domains were capable of neutralization.

Multiple approaches were undertaken to better characterize the epitopes of these antibodies. Using negative-stain electron microscopy (nsEM), images of sufficient quality to enable three dimensional reconstructions of Fab:spike protein complexes for 5 of the Fabs were collected: 3 RBD binders (Ab104, Ab138, and Ab169), and two NTD binders (Ab130 and Ab89). Although the individual antibodies have unique epitopes exhibiting different orientations of the Fab relative to the spike protein, similarities and overlaps were observed (Fig. S5). We also employed hydrogen-deuterium (H/D) exchange followed by mass spectrometry (Table S3) to obtain epitope information for antibodies not observable by nsEM, and to gain finer epitope sequence detail for several antibodies. Consistent with nsEM experiments, peptides exhibiting protection from exchange reside within the expected structural regions based on the nsEM imaging for those antibodies characterized by both methods. Epitope information was also obtained for an additional five RBD binders, three NTD binders, and three antibodies where protection from H/D exchange was not localized to a single domain (Ab82) or S2 binders (Ab127 and Ab164).

*Figures S1-S7*

#### Fig. S1. Screening and sequence analysis.

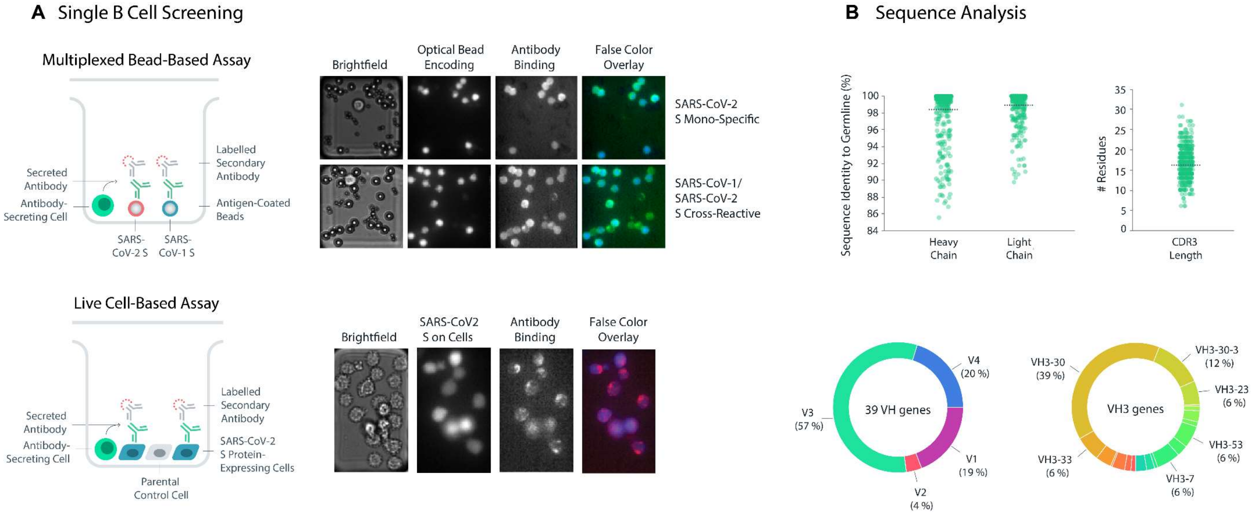

**Fig. S1. Screening and sequence analysis.** (A) (Left) Representation of multiplexed bead-based and live cell-based screening assays. (Right) Microscope images of a single selected chamber in the two assay types. The intensity of the optical beads in the multiplexed assays represents the two different antigen-coated beads, with examples of mono-specific and cross-reactive antibody binding. In the live-cell assay, SARS-CoV-2 spike protein-expressing cells are visualized by a passive dye and distinguishable from undyed non-transfected negative control cells. Positive binding is detected by fluorescence of labeled secondary antibodies. (B) Sequence analysis of the 440 unique high-confidence paired-chain antibodies. Sequence profiles of antibodies showing distributions of sequence identity to germline for heavy and light chains, CDR3 length, and VH gene usage.

### Fig. S2. Recombinant expression, biophysical characterization, and binding kinetics

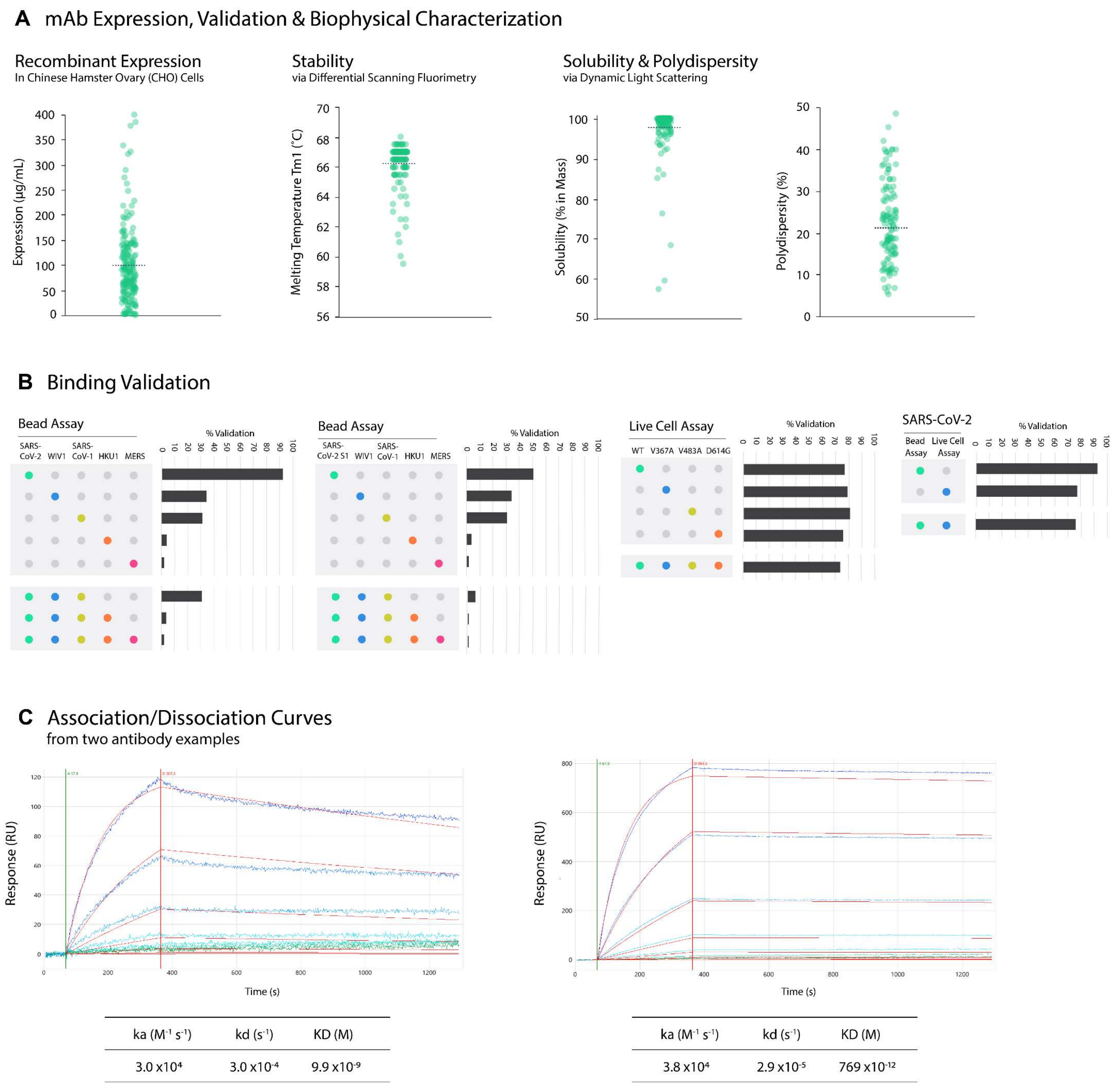

**Fig. S2. Recombinant expression, biophysical characterization, and binding kinetics.** (A) In vitro expression profile of recombinant antibodies with melting temperature, solubility, and polydispersity distributions. (B) Binding validation of recombinantly expressed antibodies. Unique antibody sequences were validated for binding using a multiplexed bead-based assay and a live cell-based assay via high-throughput flow cytometry. Antibodies were incubated with the antigen-conjugated multiplexed beads or antigen-expressed cells and positive binding was detected using a fluorescent-labeled anti-human IgG secondary antibody. The percent validation of antibodies across antigens and between assays was analyzed using AbCellera's Celium™ bioinformatics and visualization software. (C) Example association and dissociation curves for selected antibodies. Association and dissociation rate constants measured by high-throughput surface plasmon resonance (SPR) capture kinetic experiments with antibodies as immobilized ligands and antigens of interest as analytes injected at 8 concentration points. mAb = monoclonal antibody.

**Fig. S3. Epitope binning heat map.**

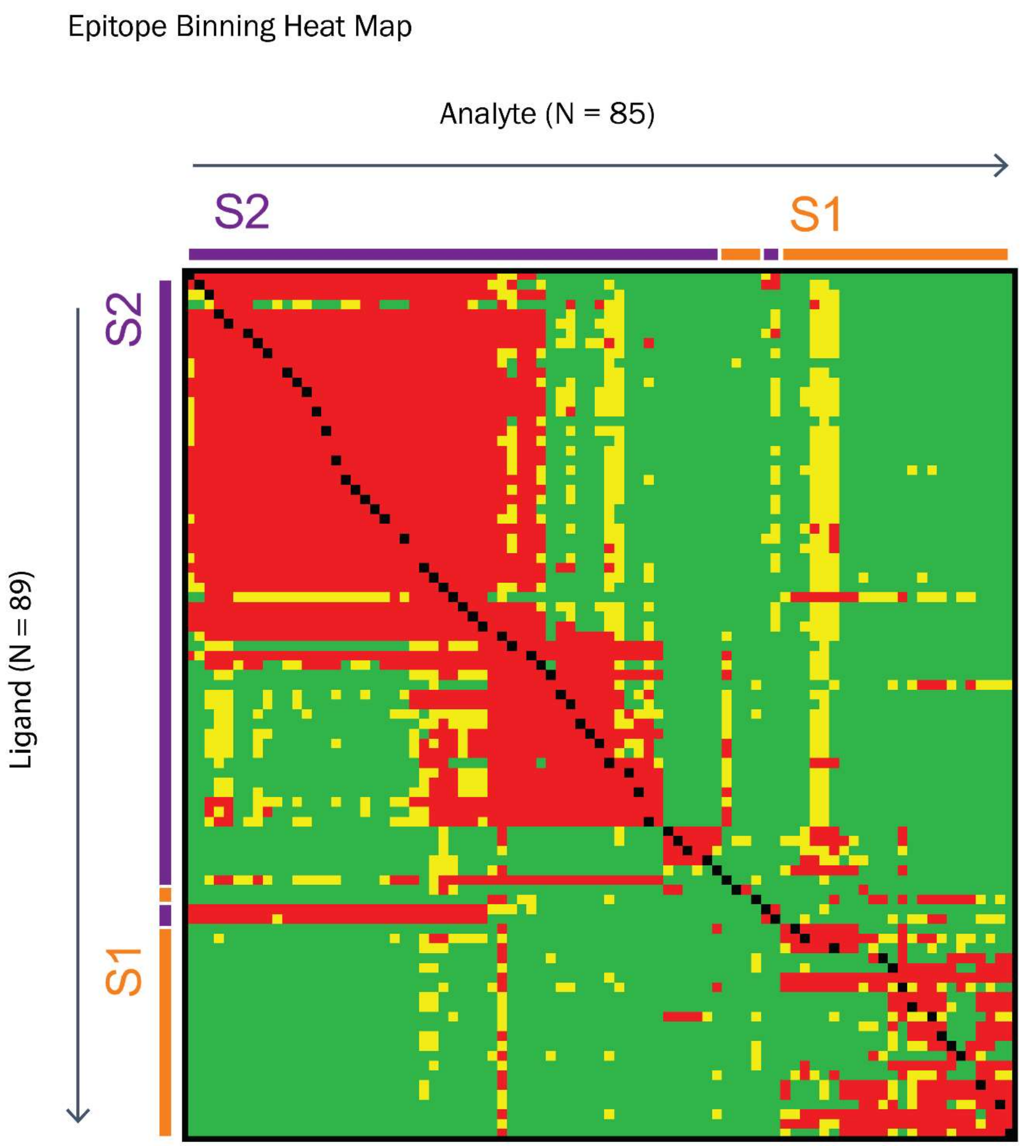

**Fig. S3. Epitope binning heat map.** The heat map shows the pairwise interaction of 85 x 89 (analyte x ligand) antibodies. Boxes in black show the self-blocking results of antibody pairs where data was available for both analyte and ligand orientations. Blocking events are highlighted in red, ambiguous binding in yellow and sandwich events in green. Binding mode as inferred by benchmark competition is shown by the purple (S2) and orange (S1) bars on the sides of the heatmap.

**Fig. S4. Epitope Binning Communities (Carterra® LSA™) for 24 selected antibodies**

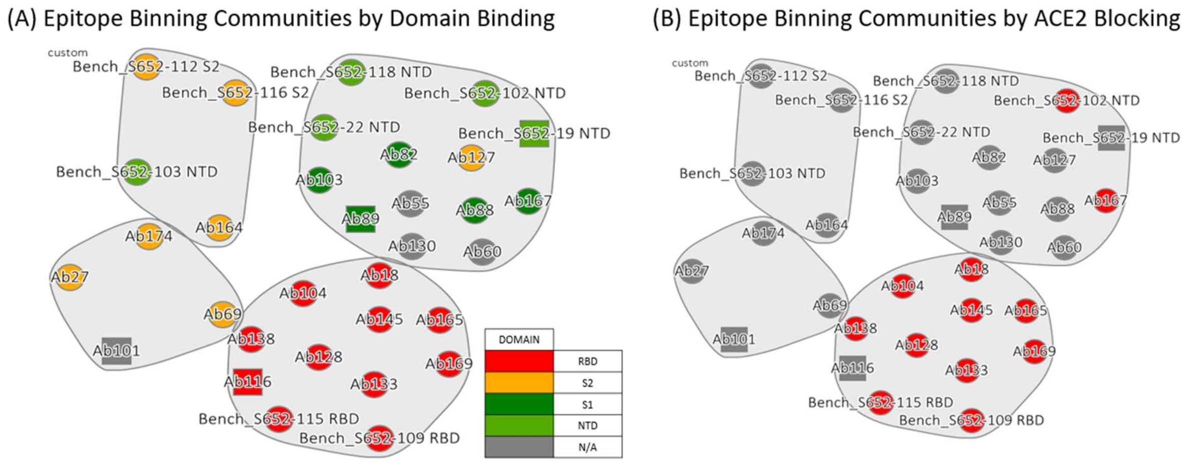

**Fig. S4. Epitope Binning Communities (Carterra® LSA™) for 24 selected antibodies.** The communities' function of the Carterra Epitope software allows for clustering of antibodies that have highly similar blocking profiles. 24 selected antibodies are grouped into 4 bins. (A) Antibodies are shown to bind the RBD domain cluster together (Red) with very little overlap between S1 domain binders (excluding RBD domain binders) and NTD benchmarks (Green) and S2 domain binders and benchmarks (Orange). (B) Antibodies annotated in red show ACE2 blocking and are dominantly found in the RBD binding community. ACE2 = angiotensin converting enzyme 2; RBD = receptor-binding domain.

**Fig. S5. Structural assessment of various antibody epitopes.**

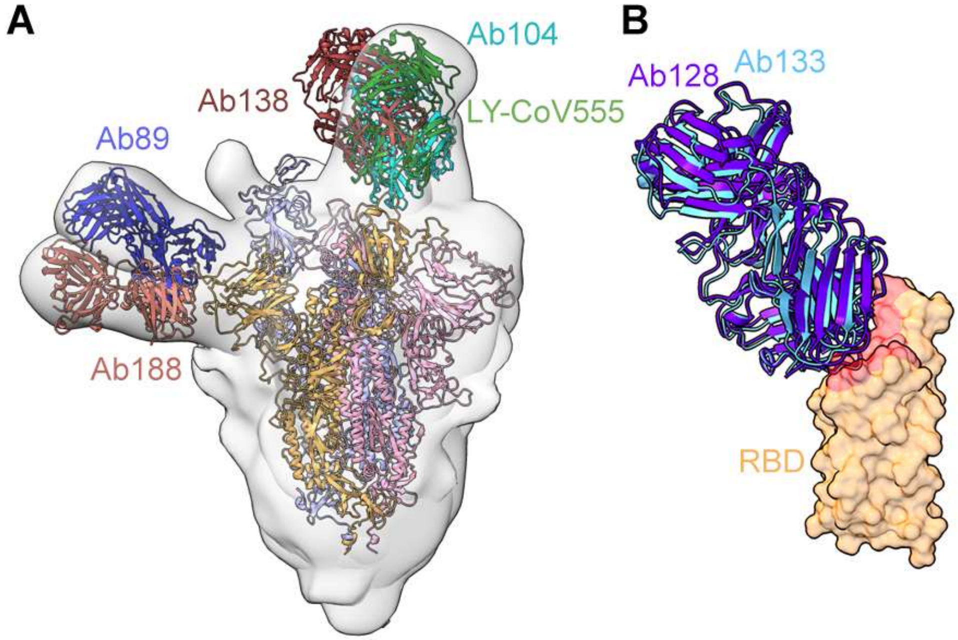

**Figs. S5. Structural assessment of various antibody epitopes.** (A) Superimposed negative-stain EM maps of spike-Fab complexes containing docked models for spike and the Fabs. (B) Superimposed RBD-Fab crystal structures for Ab128 and Ab133. The ACE2-binding surface is shown in red (PDB ID: 6M0J).(*56*) ACE2 = angiotensin converting enzyme 2; EM = electron microscopy; RBD = receptor-binding domain.

**Fig. S6. Biolayer interferometry measurements of Fab-spike binding kinetics**

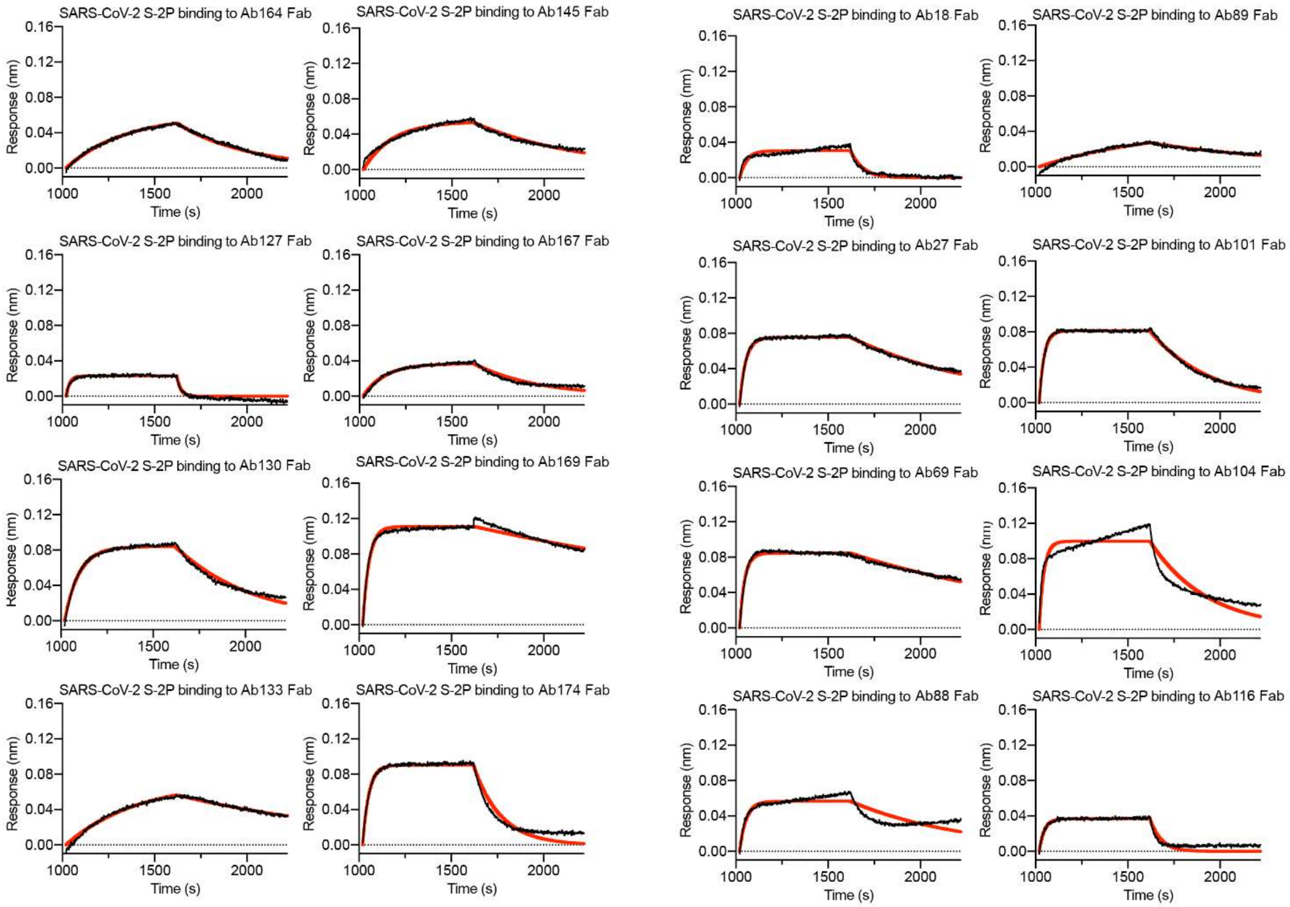

**Fig. S6. Biolayer interferometry measurements of Fab-spike binding kinetics.** Biolayer interferometry sensorgram showing binding to spike protein by the selected mAb Fabs. The data are shown as black lines and the best fit of the data to a 1:1 binding model is shown in red.

**Fig. S7. Cryo-EM structure validation**

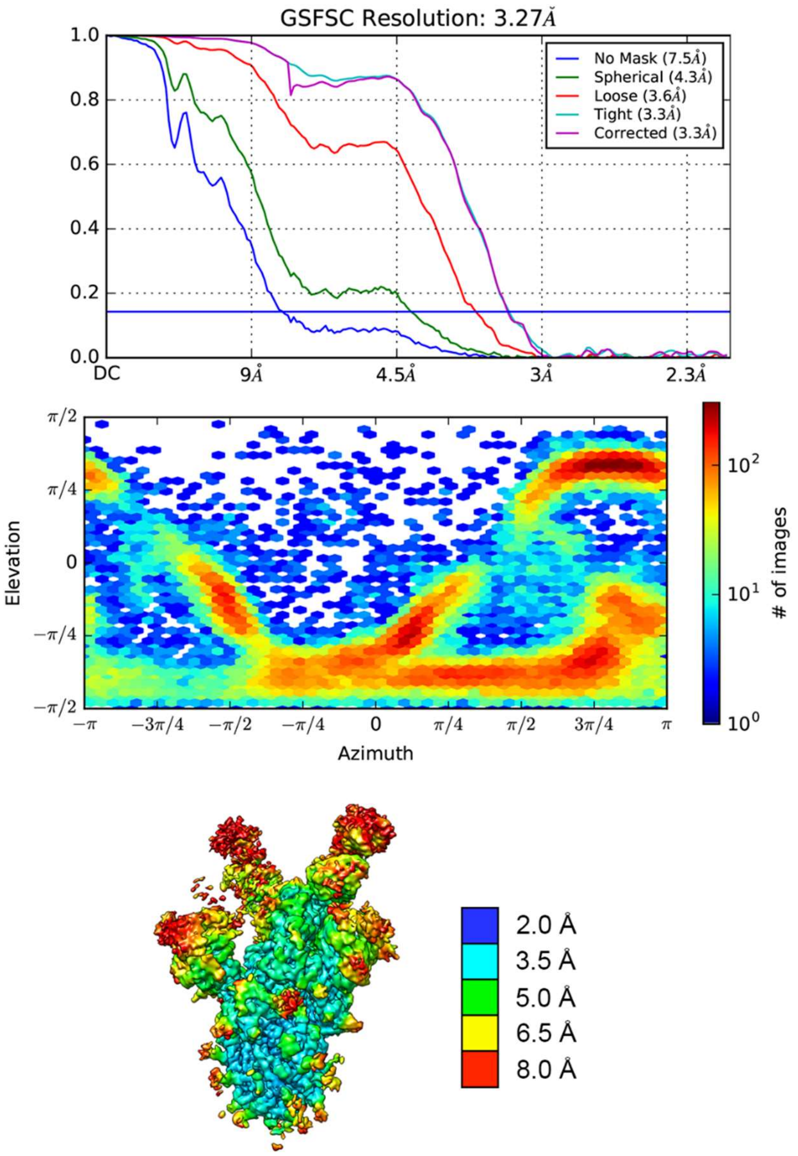

**Fig. S7. Cryo-EM structure validation**. FSC curves (Top), viewing direction distribution (Middle) and map colored by local resolution (Bottom) for the HexaPro+LY-CoV555 cryo-EM map. EM = electron microscopy.

*Tables S1-S8*

**Table S1. Selected Ab info.**

| **mAb** | V-GENE (Heavy Chain) | Identity to Germline (Heavy Chain) | V-GENE (Light Chain) | Identity to Germline (Light Chain) | **ACE2 Blocking?** |
|---|---|---|---|---|---|
| Ab18 | IGHV3-53 | 100 | IGKV1-6 | 100 | YES |
| Ab27 | IGHV3-30-3 | 100 | IGKV3-20 | 99.7 | NO |
| Ab55 | IGHV3-49 | 100 | IGKV1-5 | 100 | NO |
| Ab60 | IGHV4-59 | 100 | IGKV3-20 | 100 | NO |
| Ab69 | IGHV3-30 | 99.3 | IGKV3-20 | 100 | NO |
| Ab82 | IGHV3-30 | 99 | IGKV1-39 | 98.6 | NO |
| Ab86 | IGHV4-39 | 100 | IGKV3-20 | 100 | NO |
| Ab88 | IGHV1-46 | 100 | IGLV3-25 | 99.3 | NO |
| Ab89 | IGHV7-4-1 | 99.7 | IGKV1-39 | 99 | NO |
| Ab100 | IGHV3-30-3 | 99.3 | IGKV3-20 | 100 | NO |
| Ab103 | IGHV1-46 | 100 | IGLV3-25 | 99.7 | NO |
| Ab104 | IGHV3-66 | 100 | IGKV1-6 | 100 | YES |
| Ab116 | IGHV3-33 | 99.7 | IGKV1-5 | 99.3 | NO |
| Ab127 | IGHV3-48 | 99.7 | IGKV3-20 | 99.7 | NO |
| Ab128 | IGHV3-53 | 99.7 | IGKV1D-12 | 99.6 | YES |
| Ab130 | IGHV4-59 | 99.7 | IGLV3-1 | 99.3 | NO |

| | | | | | |
|---|---|---|---|---|---|
| Ab133 | IGHV3-53 | 99.7 | IGKV1-33 | 100 | YES |
| Ab138 | IGHV3-66 | 99.7 | IGKV1-9 | 99.7 | YES |
| Ab145 | IGHV2-5 | 99.7 | IGLV2-14 | 99.7 | YES |
| Ab164 | IGHV3-15 | 99.7 | IGKV1-5 | 100 | NO |
| Ab165 | IGHV3-53 | 99.7 | IGKV1-39 | 96.2 | YES |
| Ab166 | IGHV1-69 | 99.7 | IGLV3-25 | 99 | NO |
| Ab167 | IGHV1-18 | 99.7 | IGKV1-39 | 99.6 | YES |
| Ab169 | IGHV1-69 | 99.7 | IGKV1-39 | 99.3 | YES |
| Ab174 | IGHV3-30 | 100 | IGLV2-14 | 99.3 | NO |

Abbreviations: ACE2 = angiotensin converting enzyme 2; mAb = monoclonal antibody.

**Table S2. Epitope Binding/Affinities.**

| mAb | Domain (Experimental determination) | $K_D$ (Mab/Spike), 25°C | $K_D$ (Mab / domain), 25°C | $K_D$ (Fab/spike) 25°C |
|---|---|---|---|---|
| Ab18 | RBD | 61 pM | 710 pM RBD | 138 nM |
| Ab27 | S2 | 16 pM | 1 nM S2 | 5 nM |
| Ab55 | NTD (HDX) | 8 nM | -- | -- |
| Ab60 | NTD (HDX) | 10 nM | -- | -- |
| Ab69 | S1 (EM) | 16 pM | 1 nM S2 | 2 nM |
| Ab82 | S1 | 17 pM | 172 nM S1 | -- |
| Ab86 | Unknown | 8 nM | -- | -- |
| Ab88 | NTD (HDX) | 21 pM | 750 nM S1 | 2.8 nM |
| Ab89 | NTD (EM) | 378 pM | 262 nM S1 | 365 nM |
| Ab101 | Unknown | 2.9 nM | -- | 9 nM |
| Ab103 | S1 | 1 nM | 943 nM S1 | -- |
| Ab104 | RBD (EM) | 110 pM | 4 nM RBD | 9 nM |
| Ab116 | RBD (HDX) | 8 nM | 9 nM S2 | 88 nM |
| Ab127 | S2 (HDX) | 56 pM | 471 nM S2 | 4000 nM |
| Ab128 | RBD (HDX) | 118 pM | 15 nM RBD | -- |
| Ab130 | NTD (EM) | 53 pM | -- | 2000 nM |
| A133 | RBD (HDX) | 53 pM | 16 nM RBD | 54 nM |
| Ab138 | RBD | 84 pM | 13 nM RBD | -- |

| | | | | |
|---|---|---|---|---|
| A145 | RBD | 49 pM | 16 nM RBD | 36 nM |
| Ab164 | S2 (HDX) | 122 pM | 6 nM S2 | 452 nM |
| Ab165 | RBD | 0.3 pM | 15 nM RBD | -- |
| Ab166 | RBD | 699 pM | 2.4 nM RBD | -- |
| Ab167 | S1 | 131 pM | 175 nM S1 | 52 nM |
| Ab169 | RBD (EM & HDX) | 24 pM | 3.5 nM RBD | 1.45 nM |
| Ab174 | S2 | 173 pM | 9 nM S2 | 29 nM |

Abbreviations: EM = electron microscopy; $K_D$ = binding affinity constant; mAb = monoclonal antibody; ND = not determined; NTD = N-terminal domain; RBD = Receptor-binding domain.

#### Table S3. HDX Data summary.

| mAb | Regions showing protection |
|---|---|
| **NTD Binders** | |
| Ab55 | 136-144 (2), 171-178 (3), 242-264 (3) |
| Ab60 | 136-144 (2),171-179 (2), 242-264 (3) |
| Ab88 | 92-102 (4), 136-144 (2), 171-179(3) 242-264(3) |
| Ab89 | 136-144 (2), 242-265 (4) |
| Ab130 | 136-144 (2) 242-265 (4) |
| **RBD Binders** | |
| Ab18 | 472-495 (6) |
| Ab104 | 467-513 (9) |
| Ab116 | 467-489 (7), |
| Ab128 | 467-490 (6), 496-513(2) |
| Ab133 | 417-421 (2), 433-444 (2), 467-513 (14) |
| Ab145 | 433-455 (4), 496-513(2) |
| Ab169 | 434-444 (1), 459-495 (9) |
| **Undetermined / other binders** | |
| Ab82 | 136-143 (2), 307-318 (2), 621-636 (3) |
| Ab127 | 980-1006 (5) 1179-1186 (1) |
| Ab16 | 960-1007 (10) |

This table lists the sequence ranges exhibiting protection; the number of peptides followed for each sequence range indicated in parentheses.
Abbreviations: mAb = monoclonal antibody; NTD = N-terminal domain; RBD = receptor-binding domain.

**Table S4. Summary of viral neutralization data.**

| mAb | Pseudovirus $IC_{50}$ (Transiently transfected ACE2; VRC) | Pseudovirus $IC_{50}$ (Stably transfected ACE2; VRC) | I.F.A. $IC_{50}$ (NYU) | nanoLuc $IC_{50}$ (UNC) | PRNT $IC_{50}$ (WA isolate; UTMB) | PRNT $IC_{50}$ (Italy isolate; UTMB) |
|---|---|---|---|---|---|---|
| Ab18 | >50 µg/mL | 23 µg/mL | 1.25 µg/mL | 9.9 µg/mL | -- | -- |
| Ab27 | >50 µg/mL | >50 µg/mL | >20 µg/mL | >100 µg/mL | -- | -- |
| Ab55 | >50 µg/mL | >50 µg/mL | 10 µg/mL | >100 µg/mL | -- | -- |
| Ab60 | >50 µg/mL | >50 µg/mL | 5 µg/mL | 10.9 µg/mL | -- | -- |
| Ab69 | >50 µg/mL | >50 µg/mL | >>40 µg/mL | >100 µg/mL | -- | -- |
| Ab82 | >50 µg/mL | >50 µg/mL | 1.25 µg/mL | 1.2 µg/mL | -- | -- |
| Ab86 | >50 µg/mL | >50 µg/mL | >20 µg/mL | >100 µg/mL | -- | -- |
| Ab88 | >50 µg/mL | >50 µg/mL | >40 µg/mL | >100 µg/mL | > 100 µg/mL | >100 µg/mL |
| Ab89 | >50 µg/mL | >50 µg/mL | 0.31 µg/mL | 0.69 µg/mL | 2.47 µg/mL | 0.52 µg/mL |
| Ab101 | >50 µg/mL | >50 µg/mL | >40 µg/mL | >100 µg/mL | > 100 µg/mL | >30 µg/mL |
| Ab103 | >50 µg/mL | >50 µg/mL | -- | -- | -- | -- |
| Ab104 | >50 µg/mL | 9.8 µg/mL | 1.25 µg/mL | 1.8 µg/mL | 11.2 µg/mL | 16 µg/mL |
| Ab116 | >50 µg/mL | >50 µg/mL | 10 µg/mL | >100 µg/mL | -- | -- |
| Ab127 | >50 µg/mL | >50 µg/mL | >40 µg/mL | >100 µg/mL | -- | -- |
| Ab128 | 6.50 µg/mL | 8.6 ug/mL | 2.5 µg/mL | 5.9 µg/mL | -- | 18 µg/mL |
| Ab130 | >50 µg/mL | >50 µg/mL | 0.63 µg/mL | 27 µg/mL | > 100 µg/mL | >30 µg/mL |
| Ab133 | 0.267 µg/mL | 0.13 ug/mL | 1.25 µg/mL | 1.0 µg/mL | 0.92 µg/mL | 2.4 µg/mL |
| Ab138 | >50 µg/mL | 19 µg/mL | 2.5 µg/mL | 23.6 µg/mL | -- | -- |

| | | | | | | |
|---|---|---|---|---|---|---|
| Ab145 | >50 µg/mL | 4.0 µg/mL | 1.25 µg/mL | 2 µg/mL | -- | -- |
| Ab164 | >50 µg/mL | >50 µg/mL | >40 µg/mL | >100 µg/mL | -- | -- |
| Ab165 | >50 µg/mL | >10 µg/mL | -- | -- | -- | -- |
| Ab166 | > 12.5 µg/mL | 0.59 ug/mL | -- | -- | -- | -- |
| Ab167 | >50 µg/mL | >50 µg/mL | 20 µg/mL | >100 µg/mL | -- | -- |
| Ab169 | 0.103 µg/mL | 0.012 ug/mL | < 0.08 µg/mL | 0.03 µg/mL | 0.02 µg/mL | 0.049 µg/mL |
| Ab174 | >50 µg/mL | >50 µg/mL | >> 40 µg/mL | >100 µg/mL | -- | -- |

Abbreviations: I.F.A = immunofluorescence assay; mAb = monoclonal antibody; PRNT = Plaque Reduction Neutralization Test; IC50 = half maximal inhibitory concentration; ND = not

**Table S5: Serum Total Human IgG Concentrations and AUC 0-6days following IV Administration of LY-CoV555 or Control IgG to Rhesus Macaques in SARS-CoV-2 Challenge Model.**

| | | Serum concentration (µg /mL) | | | | |
|---|---|---|---|---|---|---|
| | | **Day 0*** | **Day 1** | **Day 3** | **Day 6** | **$AUC_{0\text{-}Day\ 6}$ (µg *hr/mL)** |
| Group 1: 50 mg/kg IgG1 control (N=4) | mean | 667 | 348 | 164 | 88 | 57500 |
| | stdev | 115 | 71 | 52 | 32 | 8360 |
| Group 2: 1 mg/kg LY-CoV555 (N=4) | mean | 15 | 13 | 10 | 8 | 1920 |
| | stdev | 3 | 3 | 2 | 1 | 356 |
| Group 3: 2.5 mg/kg LY-CoV555 (N=4) | mean | 38 | 30 | 21 | 15 | 4310 |
| | stdev | 14 | 11 | 6 | 3 | 1380 |
| Group 4: 15 mg/kg LY-CoV555 (N=3) | mean | 276 | 215 | 145 | 98 | 30900 |
| | stdev | 37 | 14 | 21 | 13 | 3190 |
| Group 5: 50 mg/kg LY-CoV555 (N=3) | mean | 679 | 539 | 376 | 258 | 77800 |
| | stdev | 101 | 61 | 47 | 78 | 12000 |

Abbreviations: $AUC_{0\text{-}Day6}$= area under the concentration time curve from Day 0 to Day 6; IgG = immunoglobulin g; IV = intravenous; N= number of animals per group; stdev = standard deviation. *Day of viral challenge.

**Table S6. Statistical analyses for impact of LY-CoV555 on viral loads in SARS-CoV-2-challenged Rhesus macaques.**

| | | BAL | | Throat Swab | | Nasal Swab | | Right Lung | | Left Lung | |
|---|---|---|---|---|---|---|---|---|---|---|---|
| | | **Genomes** | **sg mRNA** | **Genomes** | **sg mRNA** | **Genomes** | **sg mRNA** | **Genomes** | **sg mRNA** | **Genomes** | **sg mRNA** |
| | | q-val | q-val | q-val | q-val | q-val | q-val | q-val | q-val | q-val | q-val |
| **Day 1** | **1 mg/kg** | 0.159 | **0.037** | 0.084 | **0.008** | 0.207 | **0.023** | | | | |
| | **2.5 mg/kg** | 0.069 | **0.038** | **0.005** | **0.002** | 0.186 | **0.010** | | | | |
| | **15 mg/kg** | **0.008** | **0.007** | **0.001** | **0.002** | 0.454 | 0.090 | | | | |
| | **50 mg/kg** | 0.258 | 0.194 | **0.002** | **0.002** | 0.210 | **0.008** | | | | |
| **Day 3** | **1 mg/kg** | **0.022** | **0.010** | 0.730 | 0.921 | 0.069 | **0.005** | | | | |
| | **2.5 mg/kg** | **0.008** | **0.008** | 0.075 | 0.921 | **0.008** | **0.002** | | | | |
| | **15 mg/kg** | **0.007** | **0.015** | 0.794 | 0.094 | **0.031** | **0.002** | | | | |
| | **50 mg/kg** | **0.019** | **0.012** | 0.814 | 0.727 | **0.005** | **0.002** | | | | |
| **Day 6** | **1 mg/kg** | 0.192 | 0.674 | 0.250 | 0.921 | **0.027** | 0.578 | 0.137 | **0.010** | 0.137 | 0.080 |
| | **2.5 mg/kg** | **0.007** | 0.317 | **0.036** | 0.921 | **0.005** | 0.418 | **0.005** | **0.004** | **0.015** | **0.028** |
| | **15 mg/kg** | **0.011** | 0.317 | **0.037** | 0.794 | **0.038** | 0.607 | **0.002** | **0.005** | **0.005** | **0.037** |
| | **50 mg/kg** | **0.028** | 0.455 | 0.054 | 0.864 | **0.002** | 0.603 | **0.005** | **0.008** | **0.006** | **0.031** |

Abbreviation: BAL = Bronchoalveolar lavage; sg mRNA = subgenomic messenger RNA. q-values in bold represent values <0.05 and indicate statistical significance.

**Table 7. Cryo-EM data collection and refinement statistics**.

| **EM data collection and reconstruction statistics** | |
|---|---|
| Protein | SARS-CoV-2 S HexaPro+LY-CoV 555 |
| EMDB | XXX |
| Microscope | FEI Titan Krios |
| Voltage (kV) | 300 |
| Detector | Gatan K3 |
| Magnification | 22,500 |
| Pixel size (Å/pix) | 1.045 |
| Frames per exposure | 30 |
| Exposure ($e^-$/Å$^2$) | 37.2 |
| Defocus range (µm) | 0.8-2.8 |
| Micrographs collected | 2,383 |
| Particles extracted/final | 269,776 / 60,155 |
| Symmetry imposed | n/a (C1) |
| Masked resolution at 0.143 FSC (Å) | 3.27 |
| **Model refinement and validation statistics** | |
| PDB | XXX |
| Composition | |
| Amino acids | 3286 |

| | |
|---|---|
| Glycans | |
| RMSD bonds (Å) | 0.005 |
| RMSD angles (º) | 0.82 |
| Mean B-factors | |
| Amino acids | 70.5 |
| Glycans | |
| Ramachandran | |
| Favored (%) | 95.4 |
| Allowed (%) | 4.6 |
| Outliers (%) | 0.00 |
| Rotamer outliers (%) | 0.04 |
| Clash score | 10.00 |
| C-beta outliers (%) | 0.07 |
| CaBLAM outliers (%) | 2.28 |
| MolProbity score | 1.84 |
| EMRinger score | 2.40 |

**Table S8: Table of crystallographic statistics**

| | Fab Ab169 + RBD | Fab Ab133 + RBD | Fab Ab128 + RBD |
|---|---|---|---|
| **Data collection** | | | |
| Space group | P2(1) | C222(1) | C2 |
| Cell dimensions a, b, c (Å) | 42.32, 280.36, 68.91 | 74.81, 260.59, 95.00 | 105.24, 74.05, 126.09 |
| Cell dimensions alpha, beta, gamma (°) | 90, 99.62, 90 | 90, 90, 90 | 90, 111.85, 90 |
| Resolution (Å) | 30-2.16 (2.29-2.16)* | 30-1.72 (1.82-1.72)* | 30-1.73 (1.83-1.73)* |
| R-merge | 0.040 (0.444) | 0.051 (0.630) | 0.033 (0.418) |
| I / sigma (I) | 6.2 (1.5) | 13.4 (3.4) | 5.9 (1.5) |
| Completeness (%) | 91.2 (82.9) | 98.8 (98.8) | 98.7 (98.4) |
| Redundancy | 3.1 (2.7) | 13.0 (13.4) | 3.4 (3.5) |
| **Refinement** | | | |
| Resolution (Å) | 30-2.16 | 30-1.72 | 30-1.73 |
| No. of reflections | 77441 | 97212 | 93105 |
| R-work (%) / R-free (%) | 20.4 / 26.3 | 19.3 / 23.9 | 18.7 / 21.8 |
| No. of non-hydrogen atoms protein / ligand / water | 9442 / 12 / 201 | 4640 / 28 / 264 | 4813 / 26 / 492 |

| | | | |
|---|---|---|---|
| B-factors<br>protein / ligand / water | 45.9 / 33.4 / 38.4 | 43.5 / 40.2 / 40.8 | 28.1 / 45.8 / 34.3 |
| Root mean squared deviations<br>bond length (Å) /<br>bond angle (°) | 0.004 / 1.21 | 0.005 / 1.19 | 0.006 / 1.24 |
| Ramachandran distribution<br>phi-psi favored (%) /<br>phi-psi allowed (%) | 95.5 / 99.8 | 97.4 / 99.7 | 98.2 / 100 |

*Values in parenthesis denote highest resolution shell
